# RpoS activates formation of *Salmonella* Typhi biofilms and drives persistence in the gall bladder

**DOI:** 10.1101/2023.10.26.564249

**Authors:** Stuti K. Desai, Yiyang Zhou, Rahul Dilawari, Andrew L. Routh, Vsevolod Popov, Linda J. Kenney

**Affiliations:** Department of Biochemistry and Molecular Biology, University of Texas Medical Branch, Galveston, TX 77555; Sealy Center for Structural Biology and Molecular Biophysics, University of Texas Medical Branch, Galveston, TX 77555; Institute for Human Infections and Immunity, University of Texas Medical Branch, Galveston, TX 77555; Department of Pathology, University of Texas Medical Branch, Galveston, TX 77555; Department of Immunology and Microbiology, Scripps Research, 10550 N. Torrey Pines Rd., La Jolla, CA 92037, USA

**Keywords:** Static immersions, Transposon directed insertion site sequencing, ClickSeq, Asymptomatic Typhoid carriage, IraP, Fimbriae, Vi polysaccharide, CsgD

## Abstract

The development of strategies for targeting the asymptomatic carriage of *Salmonella* Typhi in chronic typhoid patients has suffered owing to our basic lack of understanding of the molecular mechanisms that enable the formation of *S.* Typhi biofilms. Traditionally, studies have relied on cholesterol-attached biofilms formed by a closely related serovar, Typhimurium, to mimic multicellular Typhi communities formed on human gallstones. In long-term infections, *S.* Typhi adopts the biofilm lifestyle to persist in vivo and survive in the carrier state, ultimately leading to the spread of infections via the fecal-oral route of transmission. In the present work, we studied *S.* Typhi biofilms directly, applied targeted as well as genome-wide genetic approaches to uncover unique biofilm components that do not conform to the CsgD-dependent pathway established in *S.* Typhimurium. We undertook a genome-wide *Tn5* mutation screen in H58, a clinically relevant multidrug resistance strain of *S.* Typhi, in gallstone-mimicking conditions. We generated New Generation Sequencing libraries based on the ClickSeq technology to identify the key regulators, IraP and RpoS, and the matrix components Sth fimbriae, Vi capsule and lipopolysaccharide. We discovered that the starvation sigma factor, RpoS, was required for the transcriptional activation of matrix-encoding genes in vitro, and for *S.* Typhi colonization in persistent infections in vivo, using a heterologous fish larval model. An *rpoS* null mutant failed to colonize the gall bladder in chronic zebrafish infections. Overall, our work uncovered a novel RpoS-driven, CsgD-independent paradigm for the formation of cholesterol-attached Typhi biofilms, and emphasized the role(s) of stress signaling pathways for adaptation in chronic infections. Our identification of the biofilm regulators in *S.* Typhi paves the way for the development of drugs against typhoid carriage, which will ultimately control the increased incidence of gall bladder cancer in typhoid carriers.

## Introduction

*Salmonella enterica* is a rod-shaped enteric bacterium that easily spreads through contaminated food or water via the fecal-oral route in poor hygiene conditions. The human-restricted serovar of *Salmonella enterica* serovar Typhi (STy), causes typhoid fever and continues to be a dangerous pathogen throughout the world[1]. The global incidence of typhoid fever in mostly children, adolescents and older adults is between 12 to 27 million and in total 116,815 succumbed to the disease in 2017[2]. In contrast, its closely related serovar Typhimurium (STm) infects diverse hosts such as humans, cattle, poultry and reptiles to cause gastroenteritis that is mostly self-limiting in healthy adults.

Upon successful invasion of intestinal epithelial cells, *Salmonella* is phagocytosed by macrophages, where it resides in a modified vacuole in a self-nourishing niche called a *<u>S</u>almonella*-<u>C</u>ontaining <u>V</u>acuole (SCV) to ultimately reach the systemic sites of liver, spleen and the gall bladder. The SsrA/B <u>T</u>wo-<u>C</u>omponent <u>R</u>egulatory <u>S</u>ystem (TCRS) is essential for activation of the *<u>S</u>almonella* <u>P</u>athogenicity Island-2 (SPI-2) regulon genes encoding a type-three secretory system and effectors that are involved in formation of the SCV[3–5]. *Salmonella* also resides encased in extracellular matrix as multicellular communities or biofilms, on intestinal epithelial cells[6], gallstones[7], tumors[8] and in the large intestine[9].

Interestingly, SsrB, a response regulator of the SsrA/B TCRS, is essential for switching on the multicellular lifestyle of *S.* Typhimurium by relieving H-NS silencing at the *csgD* promoter[10, 11]. CsgD is the master regulator of STm biofilms, and it activates the transcription of extracellular matrix components including curli fimbriae, cellulose, BapA and O-Antigen[12–16]. Moreover, the SsrB-CsgD regulatory pathway drives STm persistence in the heterologous host *Caenorhabditis elegans,* by enabling the formation of biofilms, which eventually promotes host life span through the p38-<u>M</u>itogen-<u>a</u>ctivated <u>P</u>rotein <u>K</u>inase (p38-MAPK) innate immunity pathway[17].

Biofilms in the gall bladder are important for maintaining the carrier state of *Salmonella* Typhi, allowing it to persist in 2 to 4% of chronic typhoid patients[18–20]. STy reservoirs in human carriers, who are characteristically asymptomatic, play a crucial role in the spread of typhoid in endemic regions, as well as in its introduction to non-endemic regions. Indeed, such long-term colonization of STy in human carriers, coupled with the rise of multi-drug resistant strains, for example, those belonging to the H58 haplotype, prevents effective control of typhoid fever [21–23]. Unfortunately, all of our understanding of the regulation of STy biofilms on gallstones has been based on the assumption that it shares a high degree of conservation with canonical biofilms formed by the closely related serovar, *S.* Typhimurium[18, 24].

Herein, we establish that the components of STy biofilms are fundamentally distinct from STm biofilms. In particular, CsgD and the STm lifestyle regulator, SsrB, are not required for formation of multicellular aggregates on surfaces coated with cholesterol that forms the major component of human gallstones. In order to identify unique components of STy biofilms, we employed a whole genome transposon mutagenesis approach, Tn-ClickSeq, and identified the starvation sigma factor RpoS as a crucial determinant of STy biofilms. We discovered that the formation of large STy aggregates was defective in the absence of *rpoS,* owing to the down-regulation of extracellular matrix components comprised of Sth fimbriae, the Vi polysaccharide and the lipopolysaccharide core.

Finally, we developed a heterologous host model, *Danio rerio*, to investigate STy lifestyles in persistent infections for visualizing the colonization in real-time by confocal microscopy and measuring the ensuing effects on host physiology. Zebrafish is a powerful vertebrate model for many human diseases owing to a high degree of conservation of immuno-signaling pathways and colonization characteristics of bacterial infections[25–28]. Previous studies have established that exposure of zebrafish larvae to *S.* Typhimurium, *Mycobacterium marinum*, *Shigella flexneri*, and *Pseudomonas aeruginosa* leads to successful pathogenesis[29–33]. *Shigella sonnei* and *S.* Typhimurium have also been recently shown to persistently colonize macrophages in zebrafish[34, 35]. We infected zebrafish larvae with STy using static immersions. Intestinal colonization was diminished, and larval survival was greater in the *rpoS* null, corroborating a crucial role of RpoS in prolonged STy survival in vivo. Ultimately, we harnessed the immense imaging potential of the zebrafish model to reveal the absolute requirement of RpoS in enabling chronic gall bladder colonization.

## Results

### *S.* Typhi biofilms employ unique components that differ from *S.* Typhimurium biofilms

*S.* Typhi (STy) forms biofilms on gallstones in the gall bladder and this ability is an important aspect of maintaining its carrier state[18–20]. It was therefore of interest to examine whether STy employed similar pathways as STm for biofilm formation. We grew the STy strain H58 under conditions that were proposed to mimic the gall bladder environment in humans[36], hereafter referred to as gallstone-mimicking conditions, and observed robust surface-attached communities after two days in cholesterol-coated tubes, as measured by a crystal violet staining assay (Fig. 1A). The requirement of such distinct physico-chemical factors to develop biofilms was not specific to H58, as three other isolates including Ty2-b, CT18 and CT117 formed comparable cholesterol-attached biomass after two days of growth (Fig. 1A). Not surprisingly, H58 failed to form the characteristic rough, dry and red (‘rdar’) morphotype as classically observed for the STm wild type strain 14028s (Supplementary Fig. 1A). This failure to develop rdar colonies has also been reported for other STy strains[37, 38]. Similarly, crystal violet staining of static biofilms formed in the standard ‘*Salmonella*-conditions’, of 30°C and low osmolality, demonstrated that H58 formed extremely poor biofilms when compared to the STm strain 14028s (Supplementary Fig. 1B). We followed the developmental course of H58 biofilm formation at two, four and six days in cholesterol-coated tubes and observed only a marginal increase in the amount of biofilms at days 4 and 6 compared to day 2 (Supplementary Fig. 1C). We therefore focused on understanding STy biofilms at day 2 in our further investigations.

**Figure 1:**
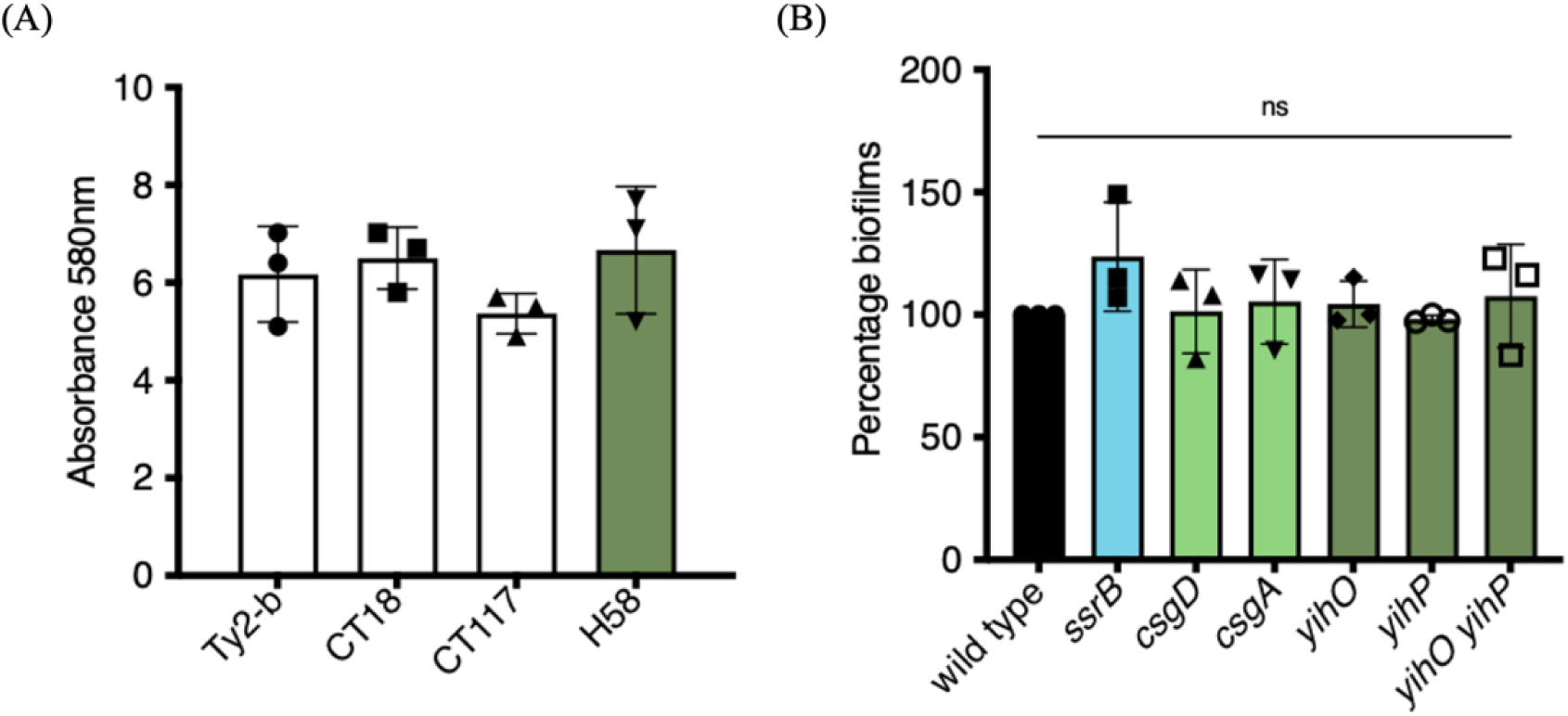
The master regulator CsgD, and the STm lifestyle regulator SsrB, are not required for the formation of cholesterol-attached STy biofilms. Crystal violet staining assays: (A) Wild type strains Ty2-b, CT18, CT117 and H58 formed robust biofilms at day 2 in gallstone-inducing conditions and (B) STy null strains deleted for the indicated STm-biofilm homologs formed similar biofilms as the H58 wild type strain at day 2. Growth medium added to cholesterol-coated Eppendorf tubes was used as the control and subtracted from all measurements. N = 3 in at least triplicates, error bars represent Mean ± SD, ns = not significant by One-way ANOVA.

The response regulator SsrB plays a dual role in regulating *S.* Typhimurium lifestyles. For virulence, SsrB∼P activates SPI-2 genes, while the unphosphorylated form drives the formation of biofilms by de-repressing *csgD*^[10, 11^ ^for^ ^a^ ^review]^. Hence, we examined whether complete deletions of *ssrB* and *csgD* affected the ability of *S.* Typhi to form cholesterol-attached biofilms. Interestingly, biofilm formation was not affected by the loss of STy homologs encoding either SsrB, the STm lifestyle regulator, or CsgD, the master regulator of STm biofilms, emphasizing the fundamental differences in mechanisms of biofilm formation in these two closely related serovars (Fig. 1B). We also investigated whether other requirements for STm biofilms: *csgA* (thin aggregative curli fimbriae), and *yihO*/*P* adhesion (O-Antigen) were required for STy aggregates to cholesterol-coated surfaces[13, 14, 39, 40]. We discovered that curli fibers and O-Antigen did not play any role in the formation of cholesterol-attached STy biofilms, strongly establishing that STy biofilms are distinct from the model STm biofilms (Fig. 1B and see Discussion).

### Tn-ClickSeq reveals the unique genetic signature of STy biofilms

Transposon-directed insertion site sequencing (TraDIS) has been immensely useful to study the genetic repertoires essential for growth in STy and *Escherichia coli*, evolution of the invasive STm lineage ST313, and biofilm formation in *E. coli* and *Pseudomonas aeruginosa*[41–45]. We applied a similar approach, Tn-ClickSeq, by combining genome-wide transposon (<u>Tn</u>) mutagenesis with ClickSeq. ClickSeq is advantageous here, since it is a fragmentation-free next-generation sequencing (NGS) library synthesis technique and is capable of generating focused NGS read data upstream of chosen target sites[46, 47]. These features greatly simplify the digital transposon display protocol and remove artifactual recombination events inherent to common NGS library preparation techniques. Using primers targeting the 3’ or 5’ ends of inserted transposons, Tn-ClickSeq can sensitively and specifically sequence the junctions of a transposon and the adjacent genomic loci of the integration site and thus identify genetic loci involved in forming cholesterol-attached biofilms in vitro (Fig. 2A and Supplementary Fig. 2A). A Tn-library in the *S.* Typhi strain H58 was kindly provided by Stephen Baker, Cambridge University, UK^[48^ ^and^ ^see^ ^Methods]^. We grew *S.* Typhi biofilms for two days using the H58-Tn library in gallstone-mimicking conditions and isolated planktonic and biofilm fractions from a pool of thirty cholesterol-coated tubes. NGS libraries were generated from genomic DNA isolated from each of these sub-populations using the Tn-ClickSeq approach[47] (Fig. 2A).

**Figure 2:**
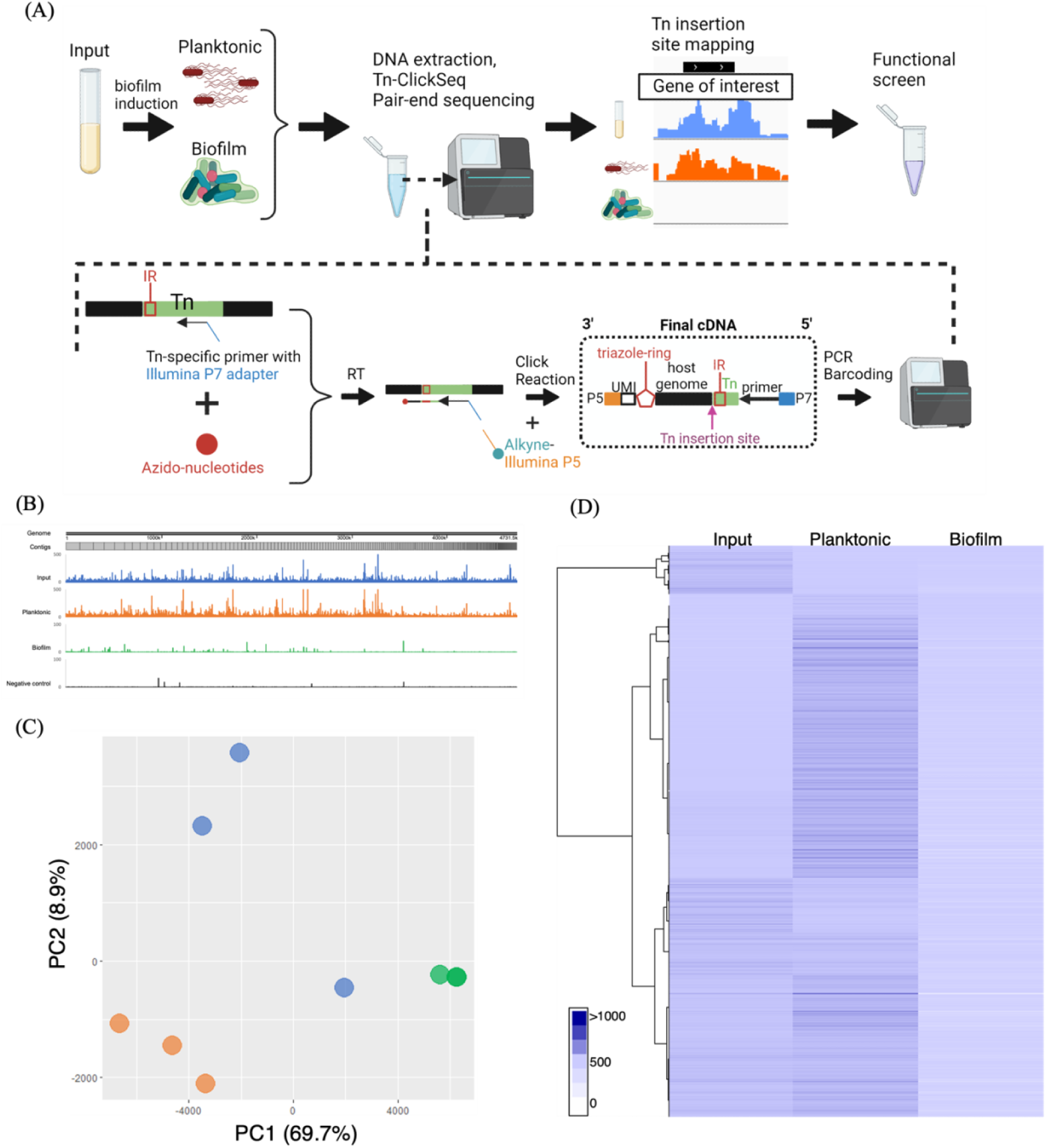
Tn-ClickSeq analysis investigates novel pathways that drive the formation of STy biofilms. (A) A general scheme depicting Tn-ClickSeq analysis, from the isolation of planktonic and biofilm fractions, DNA extraction and the generation of Tn-ClickSeq libraries using a reverse transcription reaction (RT) with a Tn-specific primer and azido-nucleotides/dNTP mixtures. The azido-terminated cDNAs were click-ligated to an alkyne-adapted illumina P5 adapter to generate Illumina libraries. After PCR and barcoding, the reads that contained partial Tn sequence and genome insertion site were pair-end sequenced and subjected to downstream analysis. (B) A linearized map of the average Tn-insertion sites from three biological replicates showed genome-wide differences in the planktonic sub-population (orange) and biofilms (green). The H58-Tn library inoculum (blue) and the H58 parent without any transposons (black, control) were used as positive and negative controls, respectively. (C) Principal component analysis showed distinct clustering of respective replicates of planktonic (orange) and biofilm (green) fractions as compared to the input library (blue) and (D) The number of transposon insertions of each gene was normalized per 1000 bp of gene length and averaged among replicates for each group to depict a hierarchical clustering of Tn-ClickSeq targets.

Since Tn-ClickSeq returns short sequence reads containing a short fragment of the 3’ or 5’ end of the inserted (known) Tn as well as a fragment of the adjacent genomic DNA, we developed a simple computational pipeline (Supplementary Fig. 2A) that identified and trimmed the Tn-derived sequences from individual reads, and then mapped the remaining fragment to the H58 genome (BioSample ID - SAMEA3110714) using a HISAT2 program[49]. The 3’ end of these Tn-mapping reads represents the exact nucleotide junction of the insertion site of the Tn in the genome. With this approach, we identified transposon insertions as 47%, 52% and 20% per million raw reads in the input, planktonic and biofilm sub-populations, respectively (Supplementary Fig. 2B, C and D). Mapping of insertion sites returned genomic locations of the inserted Tn in each dataset, as well as the frequency of these inserts within each original sample. After controlling for PCR duplication using unique molecular identifiers (UMIs) included in the ‘Click-Adaptor’[46], we assigned insertion indices by dividing the number of Tn-insertions per gene with 1 Kbp of gene length (TnClickSeq insertion indices.xlsx). With this approach, gene insertion frequencies in each condition revealed clear differences in the planktonic and biofilm fractions, indicating unique genetic components driving H58 lifestyles in gallstone-mimicking conditions (Fig. 2B). Principal Component Analysis (Fig. 2C) and hierarchical clustering analysis (Fig. 2D) returned distinct clusters of genome insertion sites of the three replicates of Tn-ClickSeq libraries obtained from the input, planktonic and biofilm sub-populations. Overall, we successfully adopted the Tn-ClickSeq approach in *S.* Typhi to generate a comprehensive view of genetic systems that determine the development of cholesterol-attached biofilms from unattached planktonic cells.

### Identifying matrix components and regulators of STy biofilms

To determine the exact mechanism(s) by which STy forms multicellular communities on cholesterol surfaces in gallstone-mimicking conditions, we compared insertion indices between the planktonic and biofilm fractions. Transposon insertions in biofilm genes likely disrupt functions resulting in the inability of such mutants to form biofilms and would be enriched in the planktonic sub-population. Interestingly, the majority of insertion indices had drastically lower values in the biofilm fraction compared to the planktonic group, these were of immediate interest for validation as STy biofilm targets (TnClickSeq insertion indices.xlsx). We focused on loci having mean insertion indices greater than 100 in the planktonic sub-population and performed Gene Ontology analysis on 1,515 such genes. We identified a significant enrichment in cell membrane components, transmembrane ion transport pathways, and other membrane-related activities (Supplementary Fig. 3). However, it was not possible to perform essentiality analysis using a standard bioinformatics pipeline, because most of these genes had insertion indices equal to zero in the biofilm fraction (see Methods and Discussion). In the absence of any *a priori* list of ‘essential’ biofilm components, we narrowed our list of 1,515 genes to the top 300 and sought to investigate some for their roles in STy biofilms. It was noteworthy that genes/operons encoding STm ‘biofilm’ homologs; *ssrB*, *csgDEFG*, *csgBAC* or *yihPO* did not appear on our list of selected Tn-ClickSeq targets. This established a strong correlation of our targeted genetic approach (Figure 1B), to our whole genome transposon mutagenesis Tn-ClickSeq approach (Fig. 2A).

We next generated precise deletions of selected loci identified by Tn-ClickSeq that might encode the structural components of STy biofilms in our gallstone-mimicking conditions: *sthC*, part of the Typhi-specific *sth* fimbrial operon[50, 51], *waaZ*, encoding an enzyme involved in the biosynthesis of the LPS core forming the outer membrane[52] and *tviD*, a part of the *tviBCDE* operon encoding the surface-exposed Vi polysaccharide (which is not encoded in STm); TviA is the regulator of the pathway[53, 54]. We performed crystal violet staining assays at day 2 and determined that the loss of *sthC*, *waaZ, tviA* and *tviD* led to a decrease in *S.* Typhi biofilms (Fig. 3A). The double mutant strains, *sthC tviD* and *waaZ sthC*, showed a similar reduction, of around 50%, as compared to the wild type parent in forming cholesterol-attached biofilms. This result indicated a possible functional redundancy or an inter-dependency in biosynthesis in the mature STy biofilms (Supplementary Fig. 4A). For example, an interplay between O-Antigen and K2 capsule synthesis has been recently described in extra-intestinal pathogenic *E. coli* for conferring serum resistance[55].

**Figure 3:**
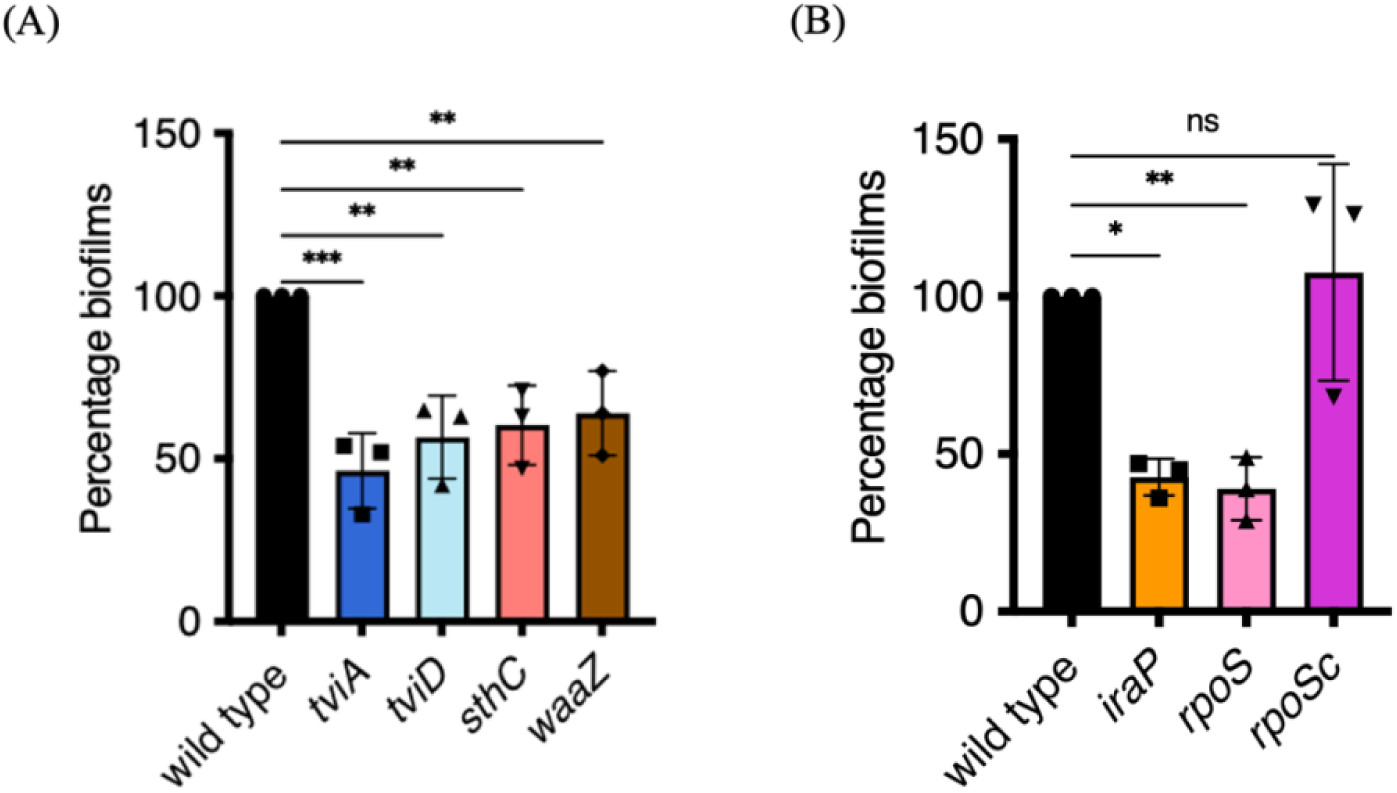
Biofilms are defective in STy null mutants identified by Tn-ClickSeq. (A) Defective biofilms formed by *tviA/tviD*, *sthC* and *waaZ* null strains suggested a role of Vi capsule, Sth fimbriae and the lipopolysaccharide core in extracellular matrix production, respectively and (B) H58 strains deleted for *iraP*, which regulates the protein stability of RpoS, and *rpoS,* which encodes the starvation sigma factor, formed less biofilms than the wild type parent. The defect in biofilm formation of the *rpoS* null strain was complemented by overexpression of *rpoS* from a plasmid *in trans*. N = 3, in at least triplicates, error bars represent Mean ± SD, in a crystal violet staining assay. Growth medium added to cholesterol-coated Eppendorf tubes was used as the control and subtracted from all measurements, ns = not significant, *p ≤ 0.05, **p ≤ 0.01 and ***p ≤ 0.001 by one-way ANOVA.

Intriguingly, the gene *iraP* rated highly on our insertion index list (insertion index = 670). It encodes an anti-adapter that inhibits the ClpXP-dependent proteolysis of the starvation master regulator RpoS[56]. We tested the biofilm-forming capabilities of an *iraP* null H58 strain and observed a substantial reduction in biofilms compared to the wild type parent at 2 days (Fig. 3B). The only known role of IraP is an indirect positive role in activating the RpoS-dependent starvation stress response in enteric bacteria[57], yet the *rpoS* insertion index was substantially lower than *iraP* (25). This was below our arbitrary threshold for selecting Tn-ClickSeq targets (insertion index > 100), thus we had excluded *rpoS* from our Gene Ontology analysis (TnClickSeq insertion indices.xlsx). It was therefore of interest to examine whether deletion of *rpoS* affected *S.* Typhi biofilms. Indeed, the *rpoS* null strain was inhibited in the formation of cholesterol-attached STy biofilms by over 50% (Fig. 3B). Over-expression of RpoS *in trans* (a kind gift from Roy Curtiss III, University of Florida), complemented the defect in biofilm formation of the *rpoS* null strain (Fig. 3B) and increased biofilms to a level similar to the wildtype. Our discovery of RpoS as an activator of *S.* Typhi biofilms was significant and highlighted the divergence between *S.* Typhi and *S.* Typhimurium biofilms. In in *S.* Typhimurium, RpoS impacts biofilms by effects on the master regulator CsgD, whereas *S.* Typhi biofilms are CsgD-independent. (Fig. 1B). Finally, we ruled out an effect of mere growth differences on biofilm formation in the STy mutants by monitoring the planktonic growth of *iraP*, *rpoS*, *sthC*, *waaZ* and *tviD* null mutants compared to the wild type H58 parent. We observed similar growth rates among all the strains, indicating that IraP, RpoS, SthC, WaaZ and TviD were necessary for the formation of *S.* Typhi surface-attached communities (Supplementary Fig. 4B).

### Cholesterol-attached aggregates of STy biofilm mutants exhibit a poor ultrastructure

Our understanding of the CsgD-dependent mechanisms of biofilm formation in *S.* Typhimurium has been achieved by screening mutant strains for their inability to exhibit the rdar morphotype and high-resolution imaging of biofilm components by fluorescence confocal microscopy[10, 37, 58–61]. Unfortunately, neither of these approaches can be applied to unequivocally establish the molecular players that drive the formation of STy biofilms on cholesterol-coated surfaces because of a high level of autofluorescence in the gallstone-mimicking conditions (see above and Discussion). We therefore grew STy biofilms in cholesterol-coated tubes for two days and obtained direct visualization of surface-attached communities by scanning electron microscopy. We observed dense aggregates of the wild type parent having a rich network of extracellular matrix that connected the cells, while the strains defective in *rpoS*, *sthC*, *waaZ* and *tviD* formed smaller STy aggregates with poor network connections (Fig. 4). Our high-resolution observations of cholesterol-attached STy communities by scanning electron microscopy consolidated our in vitro measurements of the sessile biomass formed by the STy strains by the standard crystal violet staining assay (Fig. 3). Overall, we demonstrated that the Tn-ClickSeq approach enabled the identification of previously unidentified players that drive the formation of atypical STy biofilms.

**Figure 4:**
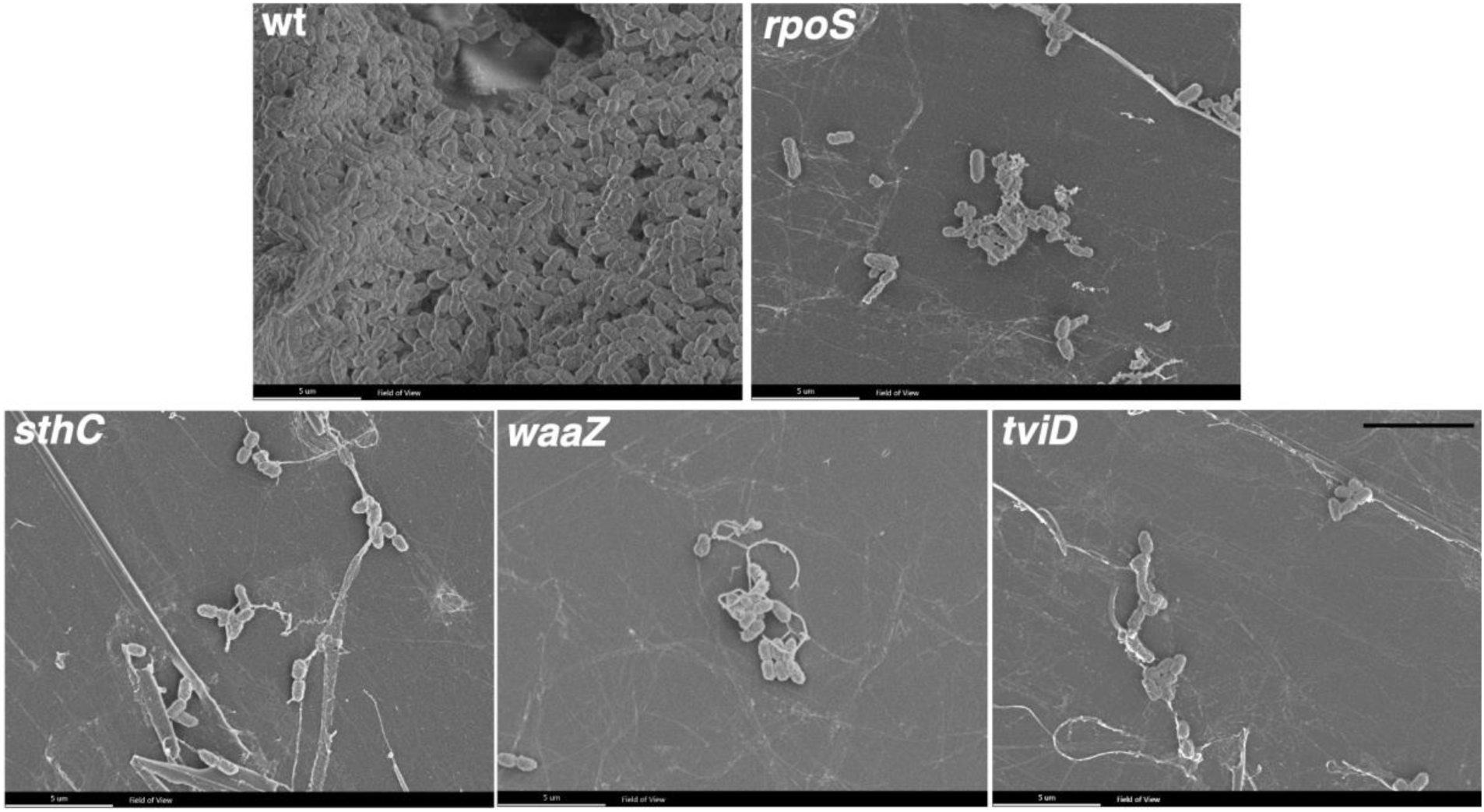
Ultra-structure visualization reveals the loss of dense aggregate formation in STy biofilm mutants. Representative scanning electron microscopy images of rich *S.* Typhi biofilms formed by the H58 (wt) and strikingly smaller-sized aggregates formed by the strains deleted of *rpoS*, *sthC*, *waaZ* or *tviD* when grown in cholesterol-coated Eppendorf tubes in gallstone-mimicking conditions. Scale bar = 5 µm.

### RpoS activates transcription of STy biofilm matrix genes

We next wanted to determine the possible role of RpoS in regulating the expression of STy biofilm components, including: SthC, WaaZ and the Vi capsule. We first investigated if the expression of *rpoS*, *sthC*, *waaZ* and *tviB* (the *tviBCDE* operon mediates biosynthesis of Vi-polysaccharide[62]), were up-regulated in biofilm-inducing conditions. We isolated planktonic and biofilm sub-populations from the wild type strain grown in cholesterol-coated tubes and measured steady state transcript levels by reverse transcription quantitative real-time PCR (RT-qPCR). Indeed, we detected an increase in the transcription of *rpoS* (∼20-fold), *sthC* (∼4-fold), *waaZ* (∼200-fold) and *tviB* (∼10-fold) in biofilms formed by the H58 parent in gallstone-mimicking conditions (Fig. 5A).

**Figure 5:**
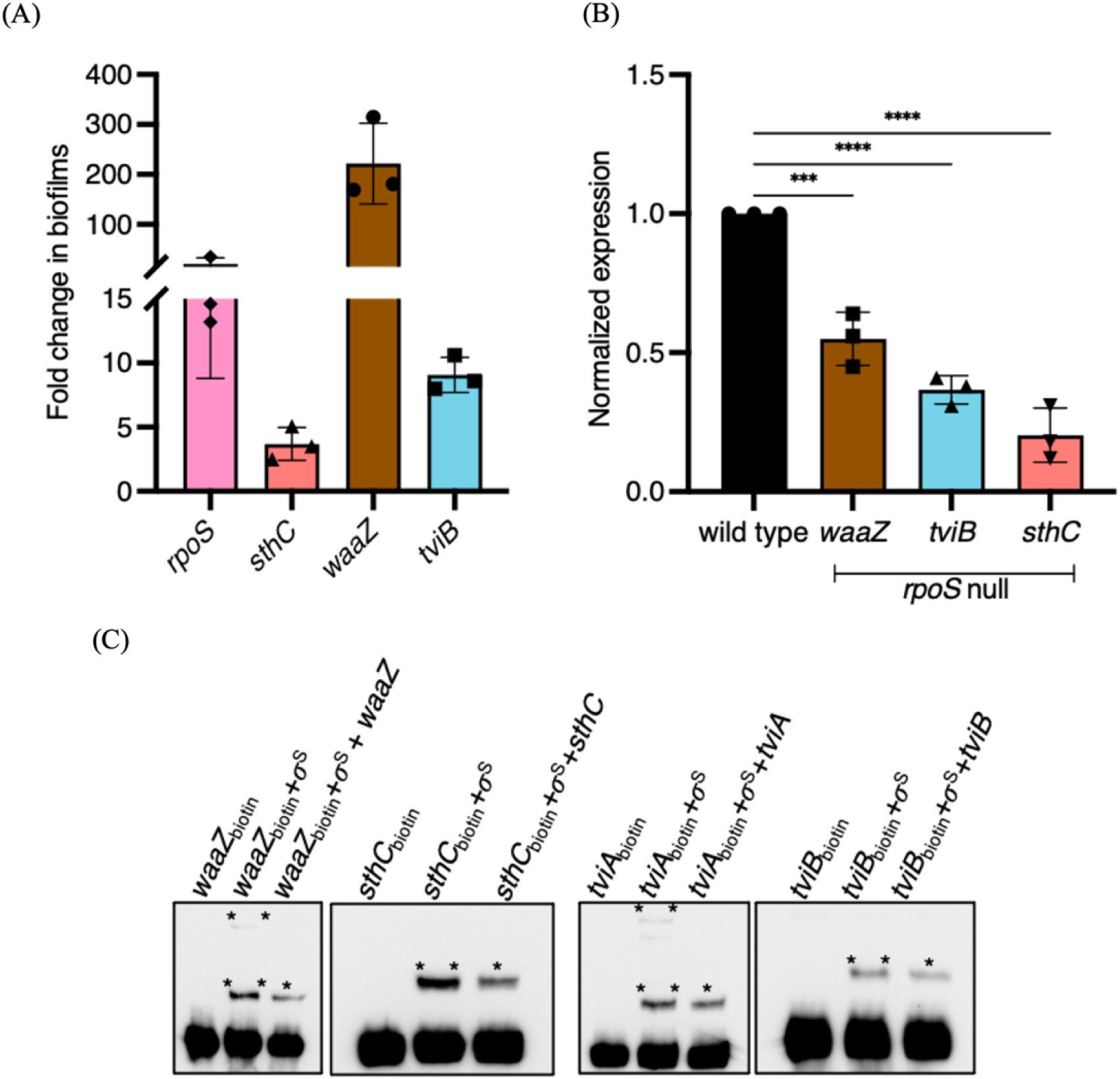
Steady-state levels of biofilm components are increased in gallstone-mimicking conditions and RpoS is required for direct transcriptional activation of biofilm matrix genes. Real time RT-qPCR analysis showed (A) a significant increase in transcription of *rpoS*, *sthC*, *waaZ* and *tviB* in the cholesterol-attached fraction compared to the planktonic fraction in the wild type H58 parent. (B) In the *rpoS* null strain, there was a significant decrease of biofilm matrix components *waaZ*, *tviD* and *sthC*. *rrsA* transcript levels were used as an internal control. N = 3, in triplicates, error bars represent Mean ± SD, ***p ≤ 0.001 and ****p ≤ 0.0001 by one-way ANOVA and (C) Electrophoretic mobility shift assays showing free biotinylated DNA, biotinylated DNA-protein complexes in the presence of RpoS (σ^S^) (**) and a decrease in the biotinylated DNA-protein complexes in the presence of excess amounts of non-biotinylated DNA (*) for *waaZ*, *sthC*, *tviA and tviB,* respectively.

In order to investigate the exact role of RpoS in driving the biofilm lifestyle, we grew the wild type parent H58 and its *rpoS* null derivative in cholesterol-coated tubes in vitro and isolated total RNA from the biofilm fractions after two days. We then compared the transcription of matrix-encoding genes *waaZ*, *sthC* and *tviB* by RT-qPCR and observed a positive effect of RpoS at each of these loci (Fig. 5B). Transcription of the matrix-encoding genes was clearly down-regulated when RpoS was absent, ∼ 60% for *waaZ*, ∼40% for *tviB* and ∼20% for *sthC* (Fig. 5B). This result established a molecular link between the stress sigma factor, RpoS, and development of STy cholesterol-attached biofilms.

Further, since the binding sites for RpoS have been previously mapped for several stress responsive genes in *E. coli*[63, 64], we analyzed the regulatory regions of *waaZ*, *sthC* and *tviA* (the regulator of the *tviBCDE* operon) in silico and found a high degree of conservation of classical RpoS recognition sequences for *waaZ* and *tviA* and a partial conservation for *sthC* (Supplementary Fig. 5A). Interestingly, binding of RpoS to the *waaZ* promoter has also been reported in *E. coli*[65]. To investigate direct binding, we amplified biotinylated fragments of these regulatory regions and tested the formation of DNA-protein complexes by electrophoretic mobility shift assays using purified His-tagged RpoS[66]. We detected RpoS binding by visualizing slower migrating DNA-protein complexes of *waaZ*, *sthC*, *tviA* and *tviB* (Fig. 5C). Binding was specific, as confirmed by a decrease in the amount of DNA-protein complexes formed in the presence of excess non-biotinylated promoter fragments as competitor DNA. Thus, RpoS directly binds and activates the formation of extracellular matrix in gallstone-mimicking conditions. RpoS binding affinity may be enhanced by the adapter protein, Crl (insertion index = 154), which was also present in our total of 1,515 Tn-ClickSeq targets (Supplementary Fig. 3; TnClickSeq insertion indices.xlsx)(see Discussion). A proposed global role of RpoS in enabling lifestyle transitions in STy is also supported by the enrichment of Tn insertions in several RpoS-regulated genes such as *gadC*, *osmB*, *bolA*, *dps*, *mscL, uspA, uspB* and *bfr*[63, 64] in the planktonic fraction as compared to the biofilm fraction (Supplementary Fig. 5B and Discussion).

### RpoS regulates long-term infection outcomes in zebrafish larvae

Since the formation of STy biofilms plays a crucial role in enabling persistence in human carriers, we investigated the role of RpoS in persistent infections in a zebrafish model. We replicated the natural mode of Typhi entry by employing static immersions of zebrafish larvae in tank water contaminated with specific doses of *S.* Typhi strains. We exposed zebrafish larvae 5 days post-fertilization (dpf) to 10^8^ cfu/ml of mCherry-expressing wild type and the isogenic *rpoS* null mutant in system water for 24 h. After 24 h, the larvae were shifted to plain system water and monitored for six days under standard conditions of zebrafish husbandry^[adapted^ ^from^ ^30]^. Control larvae were exposed to an equal volume of PBS. We discovered that *S.* Typhi colonization of infected larvae was reduced when RpoS was absent, as evident from both bacterial load measurements (Fig. 6A), as well as real-time visualizations at 2, 4 and 6 dpi using confocal fluorescence microscopy (Fig. 6B). At longer times of 4 and 6 dpi, wild type *S.* Typhi clearly persisted in the intestine (Fig. 6A, B), as also observed in a mouse model of STm persistence[67]. Finally, persistence was strongly correlated with pathogenesis outcomes, as larvae infected with the *rpoS* null strain survived infections at 6 dpi in significantly greater numbers than the wild type (Fig. 6C). The percentage survival of larvae infected with an *rpoS* mutant was similar to the uninfected PBS control from 1 to 6 dpi, strongly emphasizing the requirement of RpoS for enabling STy survival in long term infections. Monitoring infected larvae beyond 6 dpi is challenging, owing to a greater requirement of live food (Paramecia), which masks pathogen-driven physiological effects^[68^ ^and^ ^see^ ^Discussion]^. In conclusion, zebrafish infections revealed a role for RpoS in chronic STy infections and for persistence in vivo.

**Figure 6:**
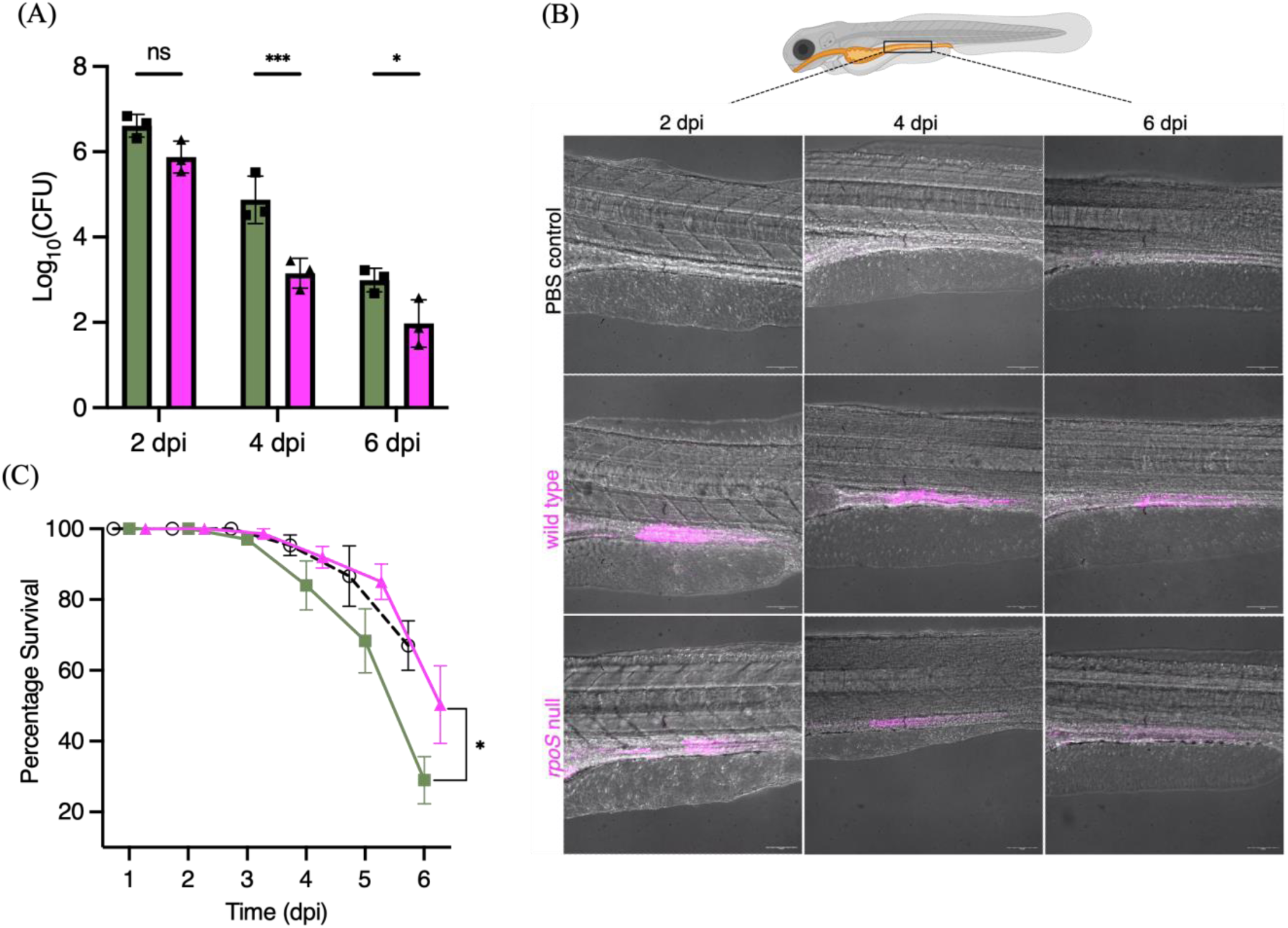
STy colonizes the intestines of chronically infected zebrafish, and RpoS is required for persistent colonization in vivo. (A) The number of STy colonies recovered from persistently infected whole larvae was drastically reduced in the *rpoS* null strain (magenta bars) compared to the wild type (green bars) at days 2, 4 and 6 post infection. Control larvae exposed to PBS were void of *Salmonella*. N = 3, five infected larvae from each group, error bars represent Mean ± SD and ns = not significant, *p ≤ 0.05 and ***p ≤ 0.001 by two-way ANOVA with Sidak’s multiple comparison tests. (B) Representative merged images (red channel and bright field) of live zebrafish larvae showing the presence of wild type mCherry-expressing *S.* Typhi in the gut. There was a stark reduction of bacteria in the infections using the mCherry-tagged *rpoS* null strain at 2, 4 and 6 dpi. No fluorescence was detected for the PBS control at all time points. The cartoon on top depicts an infected zebrafish larva with the highlighted intestinal region and was adapted from BioRender. 20x magnification, Scale bar = 100 µm and (C) Larvae infected with the *rpoS* null strain (magenta line) survived longer and showed a significant increase in the percentage of survival at 6 dpi compared to the wild type H58 (green line). The survival rate of the PBS control is shown as a dotted black line. N = 3 with 30 to 60 larvae in each group, error bars represent Mean ± SD, *p ≤ 0.05 by two-way ANOVA with Sidak’s multiple comparison tests.

### RpoS is essential for STy to persist in the gall bladder

We were then able to address the ultimate question: was RpoS necessary to colonize the hepatobiliary system? This is of importance because adaptation of STy to the gall bladder niche forms the basis for long-term transmission in humans, and increases the risks of developing hepatobiliary carcinomas. We infected zebrafish larvae with mCherry-tagged wild type H58 and *rpoS* null strains via static immersions at 5 dpf and performed whole-mount immunohistochemistry against mCherry at 6 dpi. We observed the gall bladder region by confocal fluorescence microscopy and detected anti-mCherry antibody staining in chronic infections of the H58 parent (Fig. 7, middle right zoomed-in image). Uninfected larvae were clear of any fluorescent signals in the hepatobiliary region and served as negative controls. Remarkably, we did not detect the presence of STy in the hepatobiliary system in infections using the *rpoS* null strain, emphasizing an essential role of RpoS in enabling persistent gall bladder colonization. The *rpoS* null strain was poorly visible in the intestine at 6 dpi, in complete agreement with results described above (Fig. 6A and B), further emphasizing the essentiality of RpoS signaling in hepatobiliary persistence. Since the Vi-polysaccharide capsule is also a component of the STy biofilm matrix (Fig. 4), we also performed whole-mount immunohistochemistry against the Vi polysaccharide at 6 dpi and detected clusters of bacteria, presumably in vivo STy communities, in the hepatobiliary region of zebrafish larvae infected with the wild type parent (Supplementary Fig. 6). Overall, this work provides an excellent tool, the zebrafish persistent infection model, for future investigations to unravel the exact nature of STy lifestyles in the gall bladder.

**Figure 7:**
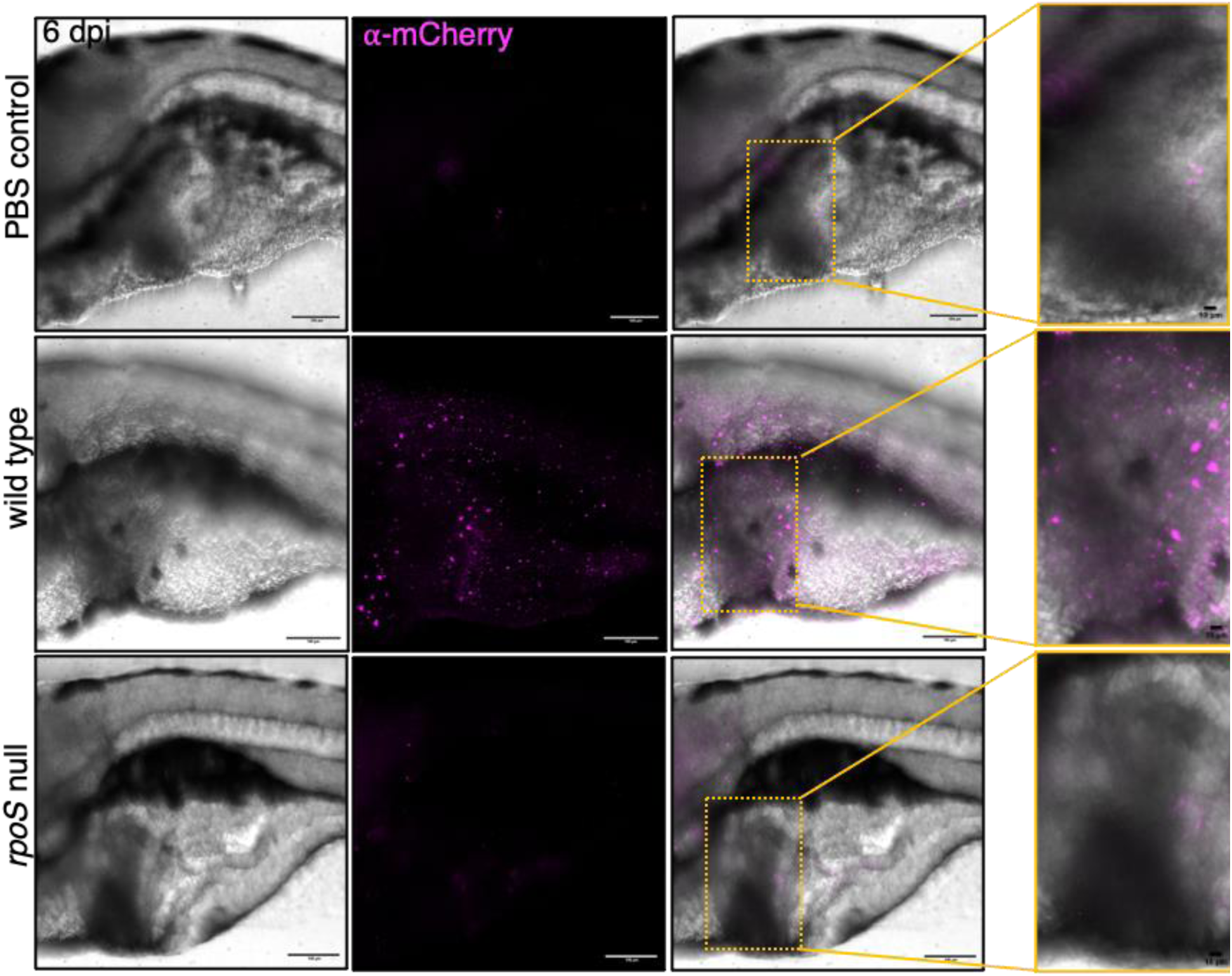
Chronic colonization in the gall bladder requires RpoS. Representative whole-mount immunohistochemistry images showing the successful detection of anti-mCherry antibody signal (magenta) in the gall bladder region of a larva infected with an mCherry-tagged wild type strain at 6 dpi (middle panel). The hepatobiliary system is highlighted by yellow rectangles in merged images and zoomed-in images show mCherry-positive fluorescent signals, validating the hepatobiliary colonization of STy in wild type infections at 6 dpi (middle right panel). In contrast, there was a drastic reduction in fluorescent signals from the gall bladder region of a larva infected with an mCherry-tagged *rpoS* null strain, with only a weak fluorescent signal as observed in the anterior intestine at 6 dpi (lower right panel, zoomed-in image). A low level of background fluorescence was also detected in the uninfected PBS control (top right panel, zoomed-in image). 20X magnification, Scale bar = 100 µm for all images except for zoomed-in images with 10 µm scale bars. N = 3 with 5 to 10 larvae analyzed in each group.

## Discussion

The dangerous transmission of *S.* Typhi from seemingly healthy, but chronically infected individuals is well documented, and the first cases in the United States of Mary Mallon, ‘Typhoid Mary’, and Mr. N in the United Kingdom, were reported in the early 1900s[69, 70]. Since then several epidemiological studies have established that persistent colonization of STy in the gall bladders of asymptomatic patients forms the basis for typhoid carriage^[reviewed^ ^in^ ^18, 24, 71]^. Despite clear evidence of the role of STy biofilms in spreading the disease, previous studies have failed to delineate any genetic mechanisms that regulate the development of gallstone biofilms in STy. This is chiefly because biofilms formed by *S.* Typhimurium have been employed as a surrogate for understanding the multicellular behavior of *S.* Typhi[72–74]. In the present work, we studied STy biofilms directly, and discovered that the pathways for biofilm formation in STy and STm were distinct. Our observations that STy failed to form biofilms in the standard laboratory conditions of low osmolality and lower temperatures (RT or 30°C) routinely employed for studying STm biofilms were in clear agreement with previous studies that observed an inability of typhoidal strains to exhibit the rdar morphotype, a hallmark of biofilm capability in the non-typhoidal *Salmonella* strains[37, 38]. More importantly, CsgD, the master regulator of STm surface-attached communities, was entirely dispensable for the formation of STy sessile aggregates on cholesterol-coated surfaces. As a result, matrix production in STy biofilms did not require curli fibers or biosynthesis of O-Antigen via the *yihO/P* system (Fig. 1B). The YihO/P permeases were previously known to transport the O-Antigen subunits for LPS biosynthesis in *E. coli* and *Salmonellae*[16, 39], but are now classified in sulfoquinovose catabolism in *E. coli*[75, 76].

### Elucidating the mechanism of STy biofilm formation

In order to identify the players that drive STy biofilms, we grew a transposon library in the STy parent strain H58 as biofilms in vitro in gallstone-mimicking conditions and developed Tn-ClickSeq analysis to map the transposon genome junctions enriched in the planktonic and biofilm sub-populations. We found Tn-genome junctions in 47% and 20% of total sequencing reads from the planktonic and biofilm fractions, respectively. These differences indicated that inactivating transposon insertions were presumably tolerated in only a low number of genes in biofilms, which correlated with their enrichment in the planktonic fraction. Also the technical challenges of isolating genomic DNA with high efficiency from STy cells attached to cholesterol-coated surfaces may affect the total sequencing yields, leading to very low insertion indices in biofilms. Nevertheless, our Tn-ClickSeq analysis generated novel insights by identifying Sth fimbriae, Vi capsule and the lipopolysaccharide core as structural components and IraP as a regulator of STy cholesterol-attached biofilms. Notably, Sth fimbriae belongs to a different fimbrial class than curli fimbriae and are not present in STm strains[50, 51]. Our comprehensive analysis of the STy biofilms also opens up exciting new directions for mapping the complete genetic signature of STy carriage.

### RpoS is a key regulator of STy biofilms

The discovery of IraP was significant, because it motivated our subsequent investigations of the downstream starvation-induced sigma factor RpoS during STy biofilm development. The reduction in biofilm formation was similar in the *iraP* and *rpoS* deletion strains (Fig. 3B), which suggested that epistatic interactions between IraP and RpoS regulated the development of STy surface-attached communities. The activation of IraP in low phosphate environments[77] provides a regulatory insight on the activation of RpoS in bi. 3Bofilm favoring gallstone-mimicking conditions. The observation that elimination of *iraP* or *rpoS* only reduced biofilms by 50% (Fig. 3B) suggests other inputs are also involved and need to be characterized. Strong candidates include the global regulators OxyR, BolA and LeuO and the TCRS RcsC/B.

Possible experimental biases in transposon library preparations may have resulted in the exclusion of *rpoS* from our list of Tn-ClickSeq targets due to its low insertion index (25). For example, the *rpoS* insertions were deterimental to survival in gallstone-mimicking conditions as shown by a higher insertion index of 65 in our input Tn-ClickSeq library compared to the planktonic sub-population. An alternative approach of serial passage of a transposon library in biofilm favoring conditions to specfically identify genetic systems regulating the development of community behavior might be informative. Nevertheless, we established that the formation of STy biofilms required RpoS and determined its role in activating the transcription of the extracellular matrix components: Sth fimbriae, Vi capsule and lipopolysaccharide (Fig. 5B). Other studies have shown that RpoS and the TCRS RcsC/B regulate Vi capsule synthesis under osmotic stress[78], and RpoS was required for the SPI-9-mediated adhesion of STy to epithelial cells[79]. The mechanisms of biofilm formation in STy are unique, in STm, the starvation response regulator RpoS activates biofilms via CsgD[80, 81], but we specifically eliminated a role for CsgD and curli in the formation of STy biofilms (Fig. 1B). We propose that RpoS directly binds to the upstream regulatory sequences of STy genes encoding the extracellular biofilm matrix components (Fig. 5C) that are AT-rich and harbor a possible extended -10 promoter element, as previously characterized for the RpoS regulon in *E. coli*[63, 64]. It is important to note here that the protein sequences of RpoS are highly conserved between *E. coli* and H58 (99% identical). A complete understanding of RpoS signaling pathways that enable the switch from the planktonic lifestyle to biofilms will require validating the precise σ^S^-recognition sequences, and hence the transcriptional start sites, for *waaZ*, *sthC*, *tviA* and *tviB*. Further understanding will also require investigating a possible co-regulatory function of Crl in stimulating the RpoS activity to enable transcriptional activation, as determined for other RpoS-responsive genes, including the *csg* operon, *adrA* and *ssrA*[66, 82–84]. Overall, the presence of other quintessential RpoS-regulated genes, such as *dps*, *bolA*, *uspA* and *osmB* in our Tn-ClickSeq dataset (Supplementary Fig. 5B ) raises an interesting question as to how RpoS coordinates stress signaling and matrix production in gallstone-mimicking conditions.

### Visualization of STy biofilms by electron microscopy

A high degree of autofluorescence and the non-specific binding of fluorescent dyes and antibodies to cholesterol surfaces prevents the visualization of mature STy biofilms using high-resolution fluorescence microscopy techniques. Although some protocols have been developed to improve the fluorescence imaging of STy aggregates in vitro and ex vivo, crucial controls that included STy strains defective in forming biofilms were lacking[85–87]. Therefore, in order to clearly visualize STy biofilms attached to cholesterol surfaces, we employed scanning electron microscopy and observed distinct aggregates of the wild type H58 parent and substantially reduced aggregate formation in the STy biofilm mutants validated from our Tn-ClickSeq analysis (Fig. 4).

### Significance of the STy-Zebrafish infection model

Our use of zebrafish larvae as a heterologous host for *S.* Typhi validated the important role of RpoS in enabling *S.* Typhi colonization in long-term infections. Reducing STy persistence by inactivating *rpoS* significantly decreased the pathogen load, prolonged host survival and most importantly, abolished hepatobiliary colonization at 6 dpi. Following STy infections beyond 6 dpi may be possible by rearing germ-free zebrafish larvae using established methods[88].

### Possible role of RpoS in the evolution of STy carriage

Finally, a strong correlation has been observed between the ability to form biofilms and the duration of STy shedding and carriage in typhoid patients from Pakistan[89]. Most recent phylogenomic analysis has proposed that the ancestral H58 haplotype originated in a chronic carrier from India and evolved to give rise to the three sub-lineages that cause a majority of typhoid infections in Asia and Africa[23]. In the light of these results, it is tempting to propose that RpoS contributes to niche adaptation in *S.* Typhi by activating the formation of biofilms in chronic carriers.

## Methods

### Bacterial strains and growth

The bacterial strains and plasmids used in this study are listed in Supplementary Table 1. *S.* Typhi strains were routinely grown in Luria-Bertani broth (LB) or Nutrient broth (NB) (BD Difco) medium with shaking at 275 rpm at 37°C in the presence of 100 µg/mL Ampicillin, 12.5 µg/mL Tetracycline, 25 µg/mL Chloramphenicol or 50 µg/mL Kanamycin (Millipore Sigma) when necessary. For the growth of STy cholesterol-attached biofilms, a modified NB medium (NB*) containing 3% w/v NB, 1.75% w/v sodium chloride, 0.25% w/v potassium chloride, 1% w/v sodium choleate and 2% w/v glucose (Millipore Sigma) was used [36]. For observing the rdar morphotype, 40 µL of overnight LB broth cultures of STm wild type strain 14028s and STy wild type strain H58 were spotted on agar plates containing 1% w/v Tryptone and 0.5% w/v Yeast Extract (LB broth without salt) supplemented with Congo Red (40 µg/mL) (Millipore Sigma) and kept at 30°C for two days. Biofilms in 96-well polystyrene plates were grown in LB broth without salt as described previously[10].

### Molecular Biology techniques

All DNA manipulation procedures were carried out according to[90] using reagents procured from Qiagen, Millipore Sigma or Invitrogen. All transformations in STy wild type strain H58 were performed by standard electroporation protocols[90]. Polymerase chain reactions (PCR) were carried out using oligonucleotides as listed in the Supplementary Table 2 following standard protocols[90].

### Strain construction

The *ssrB* null mutation in H58 strain was generated by transducing the *ssrB::kan* allele from the STm strain DW85 using standard P22 transduction protocols[91]. Other gene deletions in the STy wild type strain H58, as listed in Supplementary Table 1, were generated by the lambda *red* homologous recombination technique as described in[92, 93]. Briefly, plasmids pKD3, pKD4 or the *TetRA* DNA were used to generate linear DNA fragments by PCR using gene-specific hybrid primers as listed in Supplementary Table 2. A 10 ml LB broth culture of H58 transformed with the plasmid pKD46 (containing 100 µg/mL Ampicillin and 20 mM Arabinose) was used to electroporate 600 ng to 1 µg of purified PCR product following the protocol as described in[92]. The cells were recovered for at least 6 h at 275 rpm/30 °C after which the cells were harvested and plated on respective selective plates. The *sthC tviD* and *waaZ sthC* double null strains were generated similarly except the Kan^R^ cassettes were first removed from the *sthC* and *waaZ* null mutant strains by transforming with the temperature-sensitive plasmid, pCP20[93]. Chromosomal deletions were confirmed by PCR using flanking primer pairs as listed in Supplementary Table 2.

### STy cholesterol-attached biofilms in gallstone-mimicking conditions

STy biofilms were routinely grown following protocols adapted from[36, 39]. Sterile 1.5 ml Eppendorf tubes containing 200 µL of 10 mg/mL cholesterol (Millipore Sigma) in diethyl ether were air dried aseptically (2-3 h) and 20 µL of STy strains grown overnight in LB/NB* was added to 180 µL NB* medium. The tubes were incubated for two days at 275 rpm at 37 °C. Cholesterol-coated tubes containing only 200 µL NB* medium served as controls in all the experiments. For monitoring the time course of biofilm formation, the cholesterol-coated tubes were further incubated for four and six days, with fresh 200 µL NB* medium replaced every two days.

### Crystal violet staining assay

The amount of STy cholesterol-attached biofilms in gallstone-mimicking conditions was estimated using crystal violet staining assays adapted from[10, 39]. The supernatant/growth medium was removed, and each tube was washed once with 400 µL of Phosphate-buffered Saline (PBS). The attached biofilms were then stained with 200 µL of 0.1 % w/v crystal violet solution (filtered using Whatman Grade 1 filter paper) for five minutes at room temperature (RT). This was followed by washing once with 400 µL PBS and addition of 200 µL absolute ethanol. Appropriate dilutions were then measured for absorbance at 595 nm using an iMark™ Microplate Absorbance Reader. Each experiment was performed in triplicates or pentuplicates.

### Generation of H58-Tn library

The *TnTMDH5deloriR6K* genome library in the *S.* Typhi wild type strain H58 (H58-Tn) was kindly provided by Dr. Stephen Baker, University of Cambridge, UK[48]. The Tn5 transposon was derived from the plasmid EZ-Tn5<R6Kgori/KAN-2> (Epicentre Biotechnologies) as described in the original reference[43]. Briefly, the plasmid was digested in a 10 μL reaction containing 2.5 μL TnTMDH5deloriR6K plasmid DNA(10ng/ mL), 1μL 10X NEB4 buffer, 5.5 μL water, 0.5 μL MspAI1 restriction enzyme and 1 μL BSA, for 2 h at 37 °C. The transposon was PCR amplified using the oligonucleotides (5’-CTGTCTCTTATACACATCTC CCT and 5’-CTGTCTCTTATACACATCTCTTC) and Pfu DNA polymerase Ultra Fusion II, (Stratagene). The PCR steps were; 95 °C/ 90 seconds, followed by 30 cycles of denaturation at 95 °C/10 sec, annealing at 58 °C/20 sec and extension at 72 °C/ 20 sec, and a final extension at 72 °C/3 min.

The Tn5 amplicons were PCR purified and phosphorylated at their 5’ ends in a 80 μL phosphorylation reaction including 70 μL purified Tn5 amplicons (approximately 50 μg/ml), 8 μL 10X T4 buffer, 1 μl ATP (75 mM, Roche) and 1 μL T4 Polynucleotide Kinase (NEB). The reaction was incubated at 37 °C/45 min followed by inactivation at 65 °C/20 min. The phosphorylated Tn5 transposons were purified with a 0.5 volume of a 1:1 mixture of phenol:choloroform and centrifuged at 14,000 g/10 min/ 4 °C. The supernatant was isolated and DNA was precipitated in a 30 μL reaction including 10 μL supernatant, 1μL 3M sodium acetate (pH7.5) and 19 μL absolute ethanol. The mixture was centrifuged at 14,000 g/10 min/4 °C, the supernatant step was discarded and the pellet was rinsed twice with 200 μL of 70 % (w/v) ethanol. The pellet was eluted in 10 μL 1X Tris-EDTA (TE) buffer (pH7.5) and stored at -20 °C.

Next, transposomes were prepared using 2 μL of above purified phosphorylated Tn5 DNA (∼70 μg/mL), 4 μL EZ-Tn5 Transposase (Epicentre Biotechnologies), 2 μL 100% glycerol and 4μL 50% v/v glycerol. The reaction was carried out for 30 min/ RT. 0.2 μL of transposomes were then mixed with 60 μL electrocompetent cells of STy H58 strain for electroporation. Electrocompetent cell preparations and electrotransformations were performed exactly as described previously[43]. Finally, the kanamycin resistant colonies were resuspended in 10 % v/v glycerol for storage at -80 °C and each batch of the H58-Tn mutant library included 10 electrotransformations used to generate approximately 3 x10^5^ mutants.

### Isolation of fractions for Tn-ClickSeq libraries

A 1 µL loop of frozen stock of the H58-Tn library was inoculated in 10 mL NB* cultures and incubated overnight at 275 rpm at 37 °C. This was the ‘input’ fraction, which was used to prepare thirty tubes of cholesterol-attached biofilms in gallstone-mimicking conditions. After two days, the culture supernatants were removed and pooled to obtain the ‘planktonic’ fraction. To isolate the ‘biofilm’ fraction, 400 µL of PBS was added to each tube followed by sonication in a XUB Digital Ultrasonic Bath (Grant Instruments) for 20 min, maximum power, no leap, at RT. The harvested biomass was pooled, and the sonication step was repeated thrice. The final pool at the end of four sonication cycles was the ‘biofilm’ fraction.

### Genomic DNA isolation for generating Tn-ClickSeq libraries

Planktonic and biofilm fractions were centrifuged at 24,000 g at 4 °C for 1.5 h in a Beckman Coulter Avanti J-26XP centrifuge. Respective supernatants were discarded, and the pellets were stored on ice. 1 ml of the input fraction was also centrifuged at 15,500 g at 4 °C for 10 min and the pellet was stored on ice. Each of the pellet fractions were then resuspended in 600 µL TE buffer (pH 8.0) with 40 µL 10% w/v sodium dodecyl sulphate, 4 µl of 20mg/mL Proteinase-K (Invitrogen) and 2 µl of 10mg/mL RNaseE (Invitrogen) and mixed well by vortexing. Samples were incubated at 37 °C for 1 h, after which an equal volume of Phenol:Chloroform mixture (pH 6.7/8.0) was added, mixed and centrifuged at 15,500 g at 4 °C for 15 min. The upper aqueous phase was added to a fresh tube and an equal volume of chloroform was added, mixed, and centrifuged at 13,000 rpm at 4 °C for 15 min. The supernatants were removed, 2.5 to 3 volumes cold absolute ethanol was added, and stored overnight at -20 °C. DNA pellets were obtained by centrifuging the samples at 15,500 g at 4 °C for 15 min, followed by a 70% ethanol wash. The pellets were air dried, and DNA was resuspended in 40 µL of nuclease-free water.

### Preparation of Tn-ClickSeq libraries

For Tn-ClickSeq, genomic DNA from the input, planktonic and biofilm fractions was reverse transcribed using a reverse transcriptase (SSIII, Invitrogen) and Azido-NTPs. A Tn-specific reverse primer (3’21-39, Supplementary Table 2) was designed to the 3’ proximal end, 21 to 39 bp of transposon, with an overhang of the reverse complementary sequence of Illumina adapter (Supplementary Figure 4, Supplementary Table 2). 500 ng DNA was mixed with 1 µL of 5 µM primer, and 1 µL of 10 mM AzNTP/dTNP mixture (AzNTP:dNTP = 1:35). This initial reaction mix was heated at 95 °C for 5 min and then cooled to 50 °C in gradual steps of 0.1 °C per second. Other reaction components for the reverse transcription reaction were then added: buffer, DTT, SSIII (as per manufacturer’s protocol), and kept in a thermocycler at 50 °C for 50 min, after which the reaction was terminated at 95 °C for 5 min. This was followed by the standard ClickSeq protocol as previously described[46, 94, 95]. Briefly, immediately after denaturing, the DNA products were purified with Solid Phase Reversible Immobilization (SPRI) beads and click-ligated with a 5’-alkyne-modified adapter including a 12-nucleotide unique molecular identifier (UMI). The click-ligated product was then purified, barcoded, amplified with 18 to 20 cycles of PCR, and analyzed by agarose gel electrophoresis. The final Tn-ClickSeq libraries were subjected to pair-end sequencing on a NexSeq 550 platform at the Genomics core, UTMB.

### Bioinformatics analysis

We designed a bioinformatics pipeline to process the hybrid reads that originate from the inserted transposon and extend into the host genome (Supplementary Figure 5). The raw paired-end FASTQ reads were first pre-processed to trim the Illumina adapter, filter low-quality reads and extract UMIs using *fastp*[96]: -a AGATCGGAAGAGC -U --umi_loc read1 --umi_len 14 --umi_prefix umi -l 30. We then used *FASTX toolkit* (http://hannonlab.cshl.edu/fastxtoolkit/index.html) to reverse complement the R2 reads for ease of downstream analyses. We filtered the reads that contained the 19 bp primer sequence (targeting the Tn) with 1 nucleotide mismatch allowance with *cutadapt*[97]: -a cctatagtgagtcgtatta -e 0.1 -O 19 -m 30 -- discard-untrimmed. To obtain the reads containing primer sequences, we further filtered reads that contained the last 10 nucleotides of the 3’ invert repeat (IR) with 0% error rate allowance with *cutadapt*: - a ctgtctctta -e 0 -O 10 -m 30 --discard-untrimmed. After trimming the IR sequence, the rest of IR-containing reads were mapped to the H58 genome (https://www.ncbi.nlm.nih.gov/datasets/genome/GCF_001051385.1/) with *hisat2*[49], and then processed with *SAMtools*[98]: view/sort/index. We de-duplexed the data to minimize PCR bias with umi_tools[99]: dedup --method=unique. Finally, the locations of the transposon insertion sites were extracted with *BEDTools*[100]: genomeCoverageBed: -3 -bg. Raw sequencing data is available at the Sequence Read Archive, Project ID, PRJNA1029173.

### Gene mapping and target analysis

The number of insertion reads at each insertion site revealed by Tn-ClickSeq were ratiometrically normalized across different samples. This was followed by the assignment of gene names and annotations within H58 (GCF_001051385.1). We normalized the number of insertions to the length of each annotated gene (per 1 Kbp length of gene) to obtain the insertion indices for each samples. Matrices of all the normalized insertion dataset were then processed with DESeq2 to identify genes enriched or depleted in each conditions and perform Principal component analysis[101]. Hierarchical clustering was conducted with Cluster 3.0 (http://bonsai.hgc.jp/~mdehoon/software/cluster/), followed by TreeView (http://jtreeview.sourceforge.net/) to build the graphic map. Gene ontology analysis was conducted with GeneOntology web server (http://geneontology.org/) with *Salmonella* Typhimurium as a reference.

### RNA isolation

Planktonic and biofilm fractions were centrifuged at 24,000 g at 4 °C for 1.5 h in a Beckman Coulter Avanti J-26XP centrifuge. Respective supernatants were discarded, and the pellets were either stored at -80 °C, or immediately processed for total RNA isolation by the Trizol method[102]. Briefly, the pellets were resuspended in 1 mL TRIzol reagent (Life Technologies) and incubated at RT for 5 min. 200 µL chloroform was added, mixed well, and incubated for 3 min at RT. The mixtures were centrifuged at 15,500 g for 15 min in cold and 500 µL isopropanol was added to the supernatants. The samples were transferred to -20 °C overnight, after which the RNA was pelleted by centrifugation at 15,500 g for 15 min in cold. This was followed by a 75% ethanol wash. The pellets were air dried, and the RNA was resuspended in 20 µL of nuclease-free water.

### RT-qPCR

1 µg of total RNA extracted from the biofilm fraction of two days old cholesterol-attached biofilms was used for a reverse transcription reaction with iScript Supermix (Bio-Rad) according to the manufacturer’s protocol. This was followed by amplifying 50 ng cDNA by real-time qPCR (RT-qPCR) using SsoFast EvaGreen Supermix (Bio-Rad) and internal primers specific for *rpoS*, *sthC*, *waaZ* and *tviB*; *rrsA* was used as a normalization control (Supplementary Table 2). The annealing temperature for all the primer pairs was 56 °C. All experiments were performed in triplicates with at least three independently isolated RNA preparations. Relative expression was determined using the 2^-ΔΔC^_T_ (Livak) method as described in[102] and plotted using the GraphPad Prism 10 software.

### Scanning electron microscopy

STy cholesterol-attached biofilms were grown in gallstone-mimicking conditions for two days and after removal of the growth medium were fixed in a primary fixative containing 2.5% w/v formaldehyde (made from paraformaldehyde), 0.1% v/v glutaraldehyde, 0.01% v/v trinitrophenol and 0.03% w/v CaCl_2_ in 0.05M sodium cacodylate buffer (pH 7.3). Samples were post-fixed in 1% w/v OsO_4_ in cacodylate buffer, dehydrated in ethanol and infiltrated with hexamethyldisilazane (HMDS) to prevent cracking during drying. After air drying, the conical parts of the Eppendorf tubes were cut into strips, mounted on SEM specimen holders (metal stubs) and sputter-coated with iridium in an Emitech K575X (Emitech, Houston, TX) sputter coater for 30 seconds at 20 mA, at the Electron Microscopy Laboratory, Department of Pathology, UTMB. The samples were examined in a JEOL JSM-6330F Scanning electron microscope at the Texas Center for Superconductivity, University of Houston, at 4 µA and 2 kV.

### Zebrafish husbandry

All the protocols used for zebrafish experiments were approved by the University of Texas Medical Branch Institutional Care and Use Committee. Adult and larval zebrafish were maintained using standard husbandry procedures[103] at our in-house satellite facility. The wild type AB line was used for lifespan analysis and Typhi load measurements and the *casper* line was used for visualizing mCherry-tagged STy strains in infected larvae by confocal fluorescence imaging. Eggs were obtained by the natural spawning method[103] and kept at 28 °C in embryo medium containing methylene blue for two days. At 2 days post-fertilization (dpf), the larvae were shifted to an embryo medium containing 25 µg/mL gentamicin for 6 h, after which they were transferred to a sterile embryo medium at 28 °C. Fresh embryo medium was replaced daily.

### Static immersions of zebrafish larvae

All infections were performed using mCherry-tagged STy strains, as listed in Supplementary table 2. For static immersions, 1 ml of an overnight LB broth culture of mCherry-tagged STy strains was inoculated in 10 ml LB broth containing 100 µg/mL ampicillin and grown for 4.5 hours at 275 rpm at 37 °C. Growth was normalized by measuring absorbance at 600 nm. The strains were harvested by centrifugation at 4,200 g for 15 min at RT and resuspended in 500 µl sterile embryo medium. Ten larvae 5 dpf were added to 8 ml embryo medium in a 6-well polystyrene plate, followed by the addition of 80 µl of harvested STy cultures (10^9^ cfu/mL). Equal infection doses across strains were verified using aliquots of the harvested STy cultures by the total viable counting method. Larvae exposed to equal volume of PBS served as controls. After 24 h, the larvae were transferred to fresh embryo medium in a 6-well plate. Fresh embryo medium was replaced daily. The larvae were fed Sparos Zebrafeed (<100 µ) once daily from 9 dpf, as adopted from a delayed initial feeding model previously described[68]. Typically, infections were performed with thirty to sixty larvae in each group.

### Bacterial load estimates from infected larvae

For enumerating the STy load at 2, 4 and 6 dpi, infected larvae were isolated and added to embryo medium containing 60 to 100 µg/mL buffered Tricaine. The anesthetized larva was transferred to a 1.5 ml Eppendorf tube containing 200 µl PBS and homogenized well with a motorized micropestle. The homogenate was serially diluted and colony forming units (cfu) were estimated by the total viable counting method on LB agar plates containing 100 µg/ml ampicillin. For each time point, five larvae were used from each group. PBS control larvae remained sterile. The experiment was repeated at least three times.

### Lifespan analysis

Infected larvae were checked daily under a dissection microscope, and the percentage survival was scored. Larvae that did not show any heart beats were considered dead.

### Confocal fluorescence imaging

Five to ten larvae from each group were withdrawn at 2, 4 and 6 dpi and anesthetized in an embryo medium containing 60 to 100 µg/mL buffered Tricaine. These were then mounted on 35 mm glass-bottomed dishes with a drop of 0.8% low melting point agarose (supplemented with 60 to 100 µg/ml buffered tricaine). Images were acquired on an Olympus SpinSR-10 Yokogawa spinning disk confocal microscope fitted with an ORCA Fusion sCMOS camera (Hamamatsu) using a 20x objective (NA 0.8, Olympus), 561 CSU (or using 640 CSU for anti-mCherry IHC, below) and a step size of 1.5 µ in the Z-dimension. Images were analyzed using Image J software.

### Whole-mount immunohistochemistry (IHC) of zebrafish larvae

To detect STy colonization in the gall bladder we performed immunofluorescence of whole zebrafish larvae at 6 dpi[104]. Briefly, AB larvae infected with wild type H58 and *rpoS* null strains were fixed in 4% paraformaldehyde in PBS (pH 7.4) overnight at 4°C, washed thrice with PBS for 5 minutes and treated with a mixture of 3% hydrogen peroxide and 0.5% potassium hydroxide for 30 mins at RT to remove pigmentation. After three PBS washes, dehydration was performed using a graded methanol/PBS series of 25%, 50%, 75% and 100% with each step of 5 mins. Larvae were then kept in 100% methanol for overnight at 4°C. Rehydration steps involved a graded methanol/PBS series of 75%, 50% and 25%, with each step of 5 mins, followed by three washes with PBS containing 0.1% Tween-20 (PBS-T). For antigen retrieval, the larvae were transferred to a Tris buffer (150 mM Tris-HCl, pH 9.0) for 15 min at 70°C, washed twice with PBS-T and thrice with milli-Q water on ice. Specimens were then incubated in 100% acetone stored at -20°C for 20 mins, followed by washes with milli-Q water and PBS-T. In case of IHC using the anti-mCherry primary antibody (abcam AB213511), an additional step of treatment with 0.3% v/v Triton-X 100 for 10min/RT was included at this step. Samples were washed twice with PBS-T for 5 min before blocking in 10% BSA in PBS-T for 2h at RT, washed with PBS-T and kept overnight in 1% BSA solution in PBS-T containing a 1:100 dilution of anti-mCherry antibody or 1:500 dilution of anti-rabbit-Vi-polysaccharide antibody (BD BioSciences) overnight at 4°C. Subsequently, four PBS-T washes of 15 min each were performed, and the larvae were incubated in a secondary antibody solution containing 1% BSA in PBS-T and 1:1000 donkey-anti-rabbit 647 (for α-mCherry primary antibody) or 1:1000 goat-anti-rabbit-Alexa Fluor 555 antibody (for α-Vi-capsule primary antibody, Invitrogen) for overnight at 4°C. Final washes included four PBS-T washes and one PBS wash at RT. The larvae were then mounted on agarose and imaged on an Olympus SpinSR-10 Yokogawa spinning disk confocal microscope as described in the section on confocal imaging.

### Purification of His-RpoS protein

His_6_-tagged RpoS protein was purified from an *E. coli* DH5α strain transformed with the plasmid pUHE21-lacI^q^::*rpoS*[66] (expresses RpoS from STm 14028s strain which is 100% identical to RpoS from STy H58) following the protocol as described in Kim et al., Front Microbiol, 2021, with the following modifications: A 30 ml culture was grown in LB broth containing 100 µg/mL ampicillin till 0.6 OD_600_ and induced with 0.05 mM isopropyl-β-D-1-galactopyranoside (IPTG) for 7 hours. After sonication cells were resuspended in a lysis buffer (Buffer A) containing 50 mM NaH_2_PO4 and 300 mM NaCl, pH 8.0. A Nickel-Nitrilotriacetic acid (Ni-NTA, GOLDBIO) column was washed with water and equilibrated in Buffer A containing 30 mM imidazole. The cell lysates were incubated in Ni-NTA resin for an hour, pelleted and washed thrice with Buffer A containing 30 mM imidazole. Protein fractions were eluted in Buffer A containing 300 mM imidazole and dialyzed using a 6 to 8K MWCO dialysis tubing at 4 °C in a buffer containing 20mM Tris-HCl, pH 8.0, 150 mM NaCl, 0.1mM EDTA, 5mM dithiothreitol and 20% v/v glycerol for 2 to 3 hours. The dialyzed protein was quantified by measuring the absorbance at 280 nm using a NanoDrop (Thermo Fisher Scientific) and aliquots were stored at -20 °C for immediate use or at -80 °C for future use.

### Electrophoretic mobility shift assay (EMSA)

Electrophoretic mobility shift assays were performed using the LightShift^TM^ Chemiluminescent EMSA Kit (Thermo Fisher Scientific) according to the manufacturer’s protocol. Primers used for amplifying the upstream regulatory regions of *waaZ*, *sthC*, *tviA* and *tviB* are described in Supplementary Table 2. Biotinylated forward primers were used for generating respective biotinylated DNA fragments by PCR. Each 20 µl binding reaction contained 1 ng DNA, 2 µL binding buffer, 1 µl 1µg/µL poly dI.dC, 1 µL 50% glycerol, 1 µL 1M KCl and 1 µL 1% w/v NP-40. 1.4 µM purified RpoS, 125 ng non-biotinylated *waaZ* or *sthC* DNA and 160 ng non-biotinylated *tviA* and *tviB* were added when appropriate.

### Statistics

The Prism 10 software was used to plot all graphs and perform statistical analyses.

## Acknowledgements

S.D. and L.J.K. thank the following for their assistance; Stephen Baker (University of Cambridge) and Tran Vu Thieu Nga for the H58 Tn library, Swaine Chen, Kurosh Mehershahi and Varnica Khetrapal for initial attempts at TraDIS at the Genome Institute of Singapore, Zhixia Ding (Department of Pathology, UTMB) and Dezhi Wang (The Texas Center for Superconductivity, University of Houston) for SEM sample preparations and image acquisitions, Hyunjin Yoon and Eunsuk Kim, Ajou University, South Korea, for the His-RpoS construct, Leslie Morgan in the LJK group for His-RpoS purification and Roy Curtiss III and Soo-Young Wanda, University of Florida, for the *rpoS* construct. S.K.D. and R.D. thank the Zebrafish International Resource Center (ZIRC) for fish lines and April Freeman (ZIRC) for advice on fish husbandry. Z.Y. and A.L.R. were supported by a subaward from U54 Center Grant, AI170855, NIH/NIAID. L.J.K. acknowledges support from UTMB Start up Funds, Texas Star Award, and Cancer prevention and Research Institute of Texas (CPRIT) Grant RP2000650.

---

**Supplementary Figure 1:**
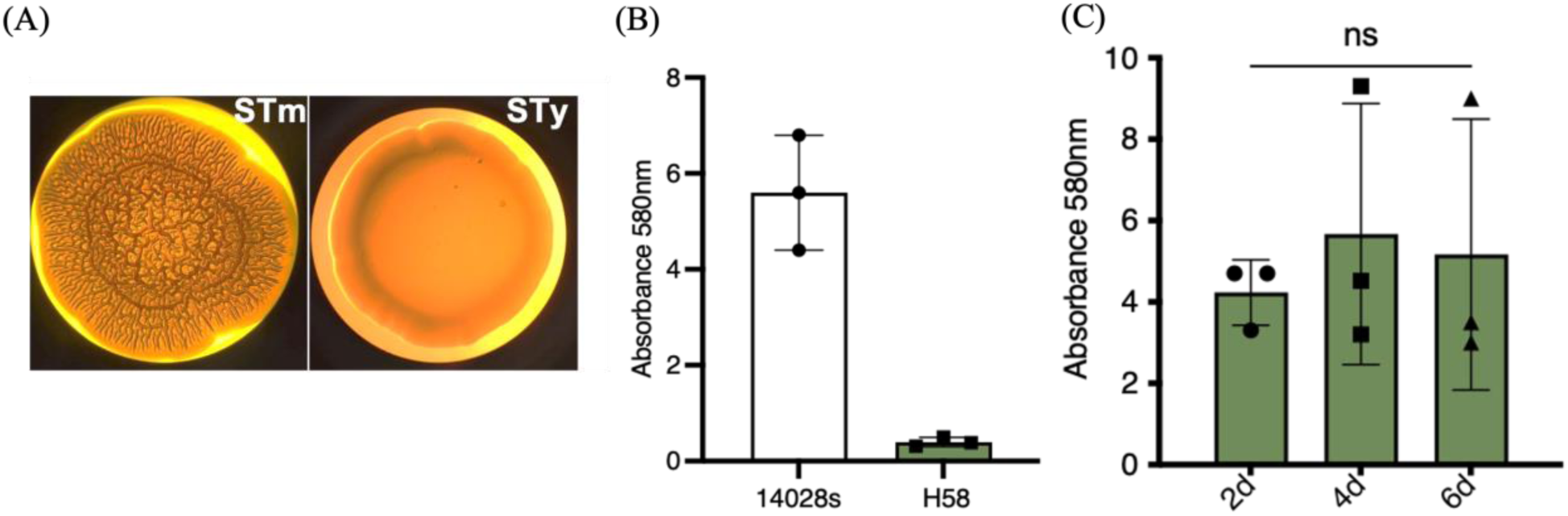
H58 forms ‘atypical’ biofilms. (A) Examination of a macrocolony of *S.* Typhimurium (STm) displaying a red, <u>d</u>ry and rough (rdar) morphotype on Luria-Bertani (LB) agar without salt medium containing the congo red dye, while *S.* Typhi (STy) are ‘smooth’ and lack the rdar morphology. (B) Wild type STy strain H58 was unable to form biofilms compared to the wild type STm strain 14028s, when grown in LB broth without salt medium in polystyrene plates at two days as determined by a crystal violet staining assay and (C) The cholesterol-attached biomass formed by wild type H58 did not increase significantly at days 4 and 6 compared to day 2, as determined by a crystal violet staining assay. Growth medium added to cholesterol-coated Eppendorf tubes was used as the control and subtracted from all measurements. N = 3, in at least triplicates, error bars represent Mean ± SD, ns = not significant by one-way ANOVA.

**Supplementary Figure 2:**
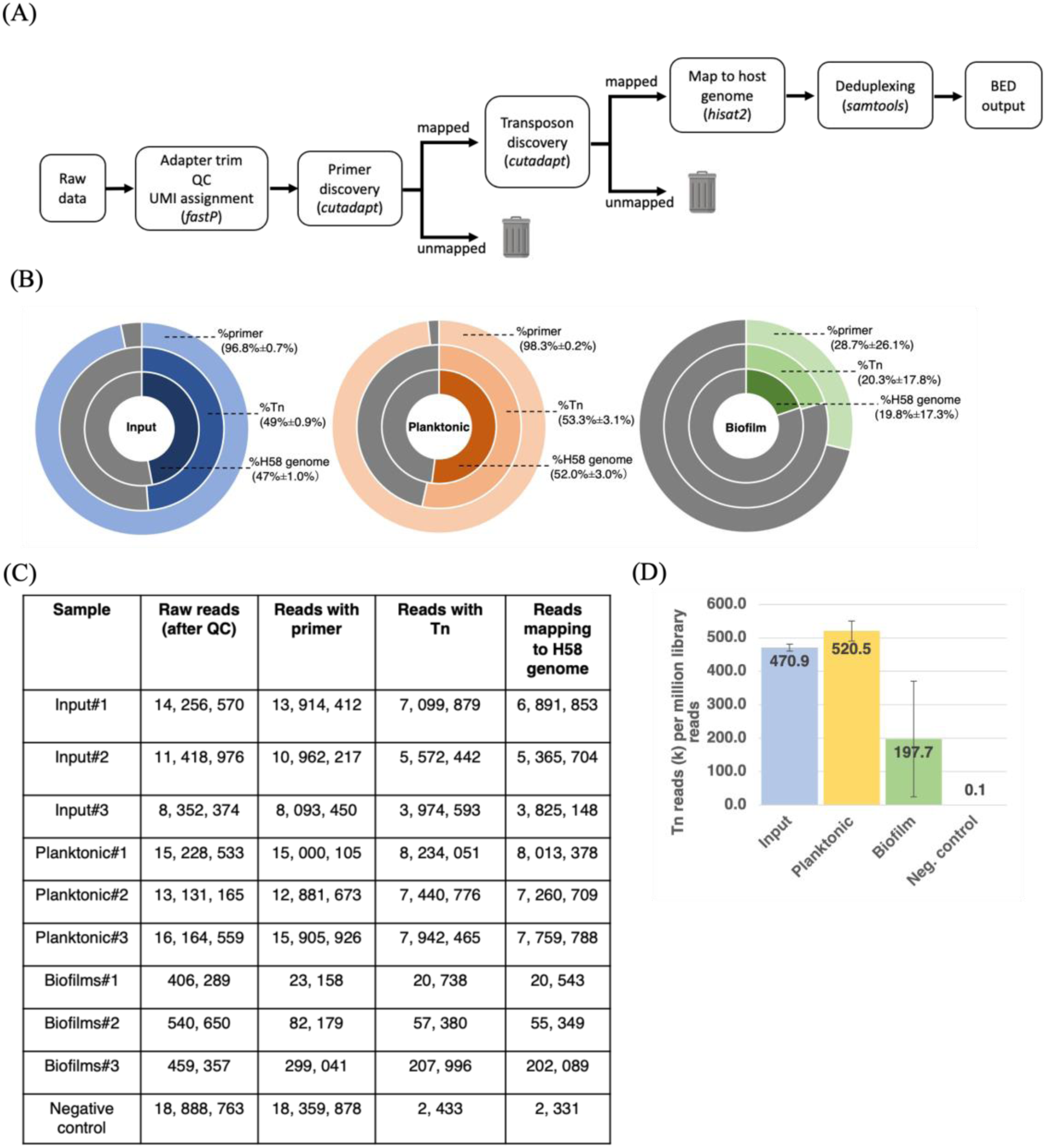
(A) A flowchart of the bioinformatics pipeline employed for Tn-ClickSeq analysis depicting the filtering of raw reads containing both primer and partial Tn sequences, the subsequent identification of genome insertion sites using the reference H58 genome and lastly de-duplexing to remove PCR biases. (B) Percentages of reads that matched the primer, transposon sequence (IR sequence) and the host genome from left to right: input, planktonic and biofilm sub-populations. (C) Sequencing yields of the respective input, planktonic and biofilm Tn-ClickSeq libraries in triplicates and of an H58 strain without any transposon insertions as a negative control and (D) Graphical representation of Tn insertion sites per one million of raw reads showing an average of 470,000 reads for the input fraction, 520,000 for the planktonic fraction and 197,000 for the biofilm fraction. A similar analysis using an H58 strain without any transposon insertions resulted in only 100 false positives.

**Supplementary Figure 3:**
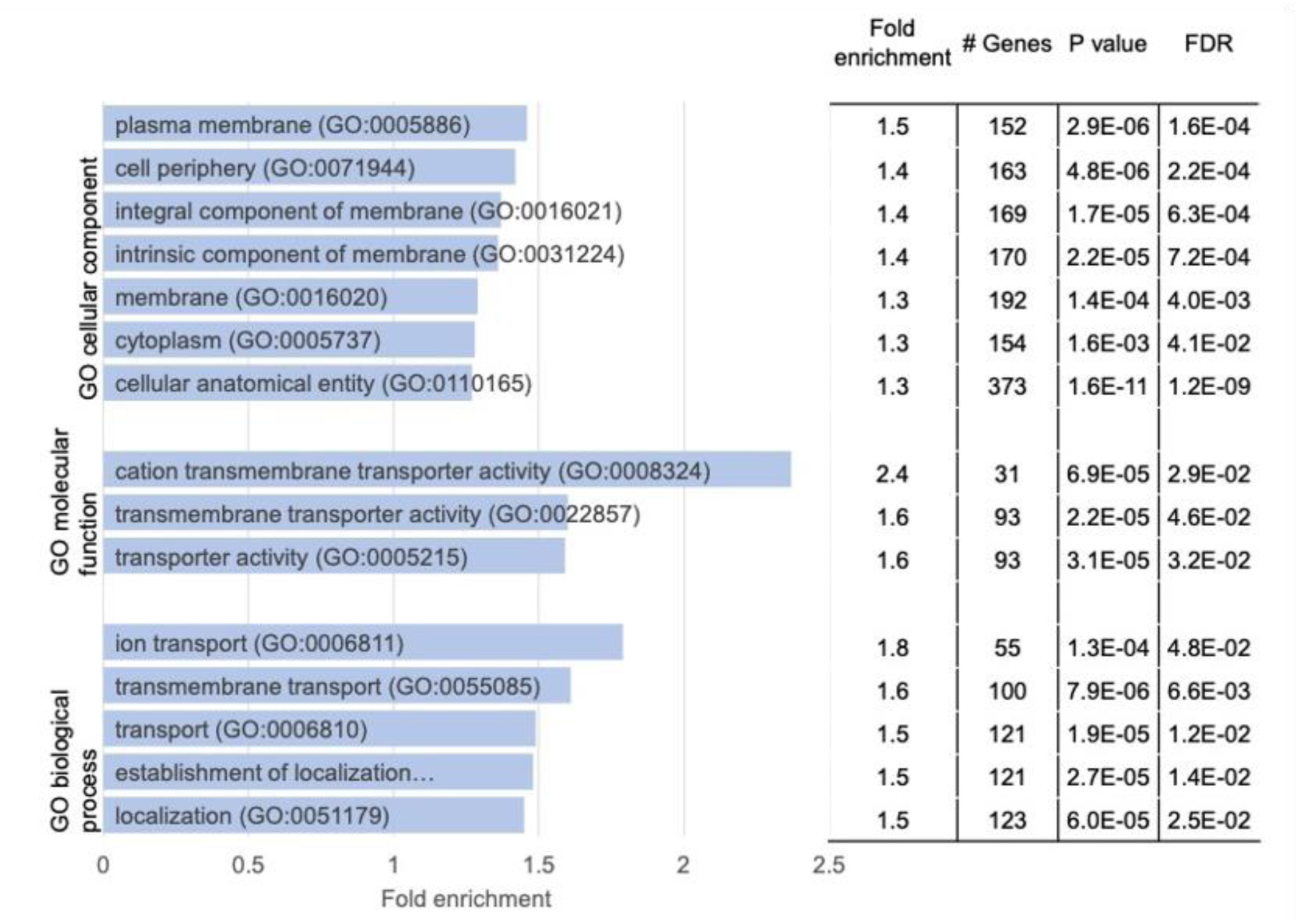
A Gene Ontology analysis of transposon insertion sites in 1515 genes that were enriched in the planktonic sub-population as compared to biofilms showed a significant enrichment of cell membrane components, transmembrane ion transport pathways and other membrane related activities.

**Supplementary Figure 4:**
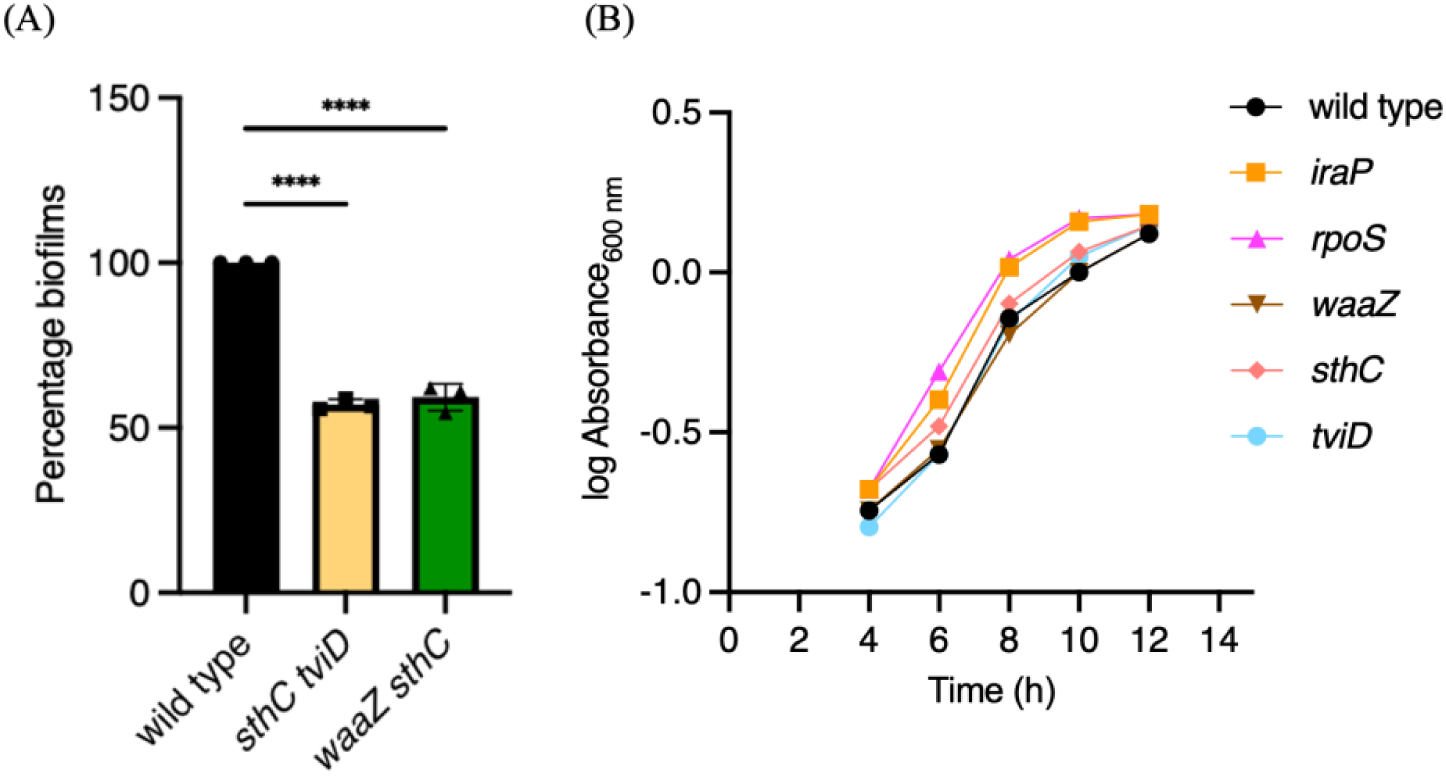
(A) Double null mutant derivatives of an H58 parent defective for Sth fimbriae and Vi-polysaccharide (*sthC tviD*), and for LPS biosynthesis and Sth fimbriae (*waaZ sthC*) showed a reduced ability to form cholesterol-attached biofilms, by around 50%, compared to the wild type parent. N = 3, in at least triplicates, error bars represent Mean ± SD, in a crystal violet staining assay. Growth medium added to cholesterol-coated Eppendorf tubes was used as the control and subtracted from all measurements, ****p ≤ 0.0001 by one-way ANOVA and (B) Biofilm mutants are not compromised in planktonic growth - Strains deleted of *iraP*, *rpoS*, *waaZ*, *sthC* or *tviD* and the wild type H58 strain were grown for 12 hours in Luria-Bertani broth at 37°C/250 rpm and the absorbance at 600 nm was measured every 2 hours.

**Supplementary Figure 5:**
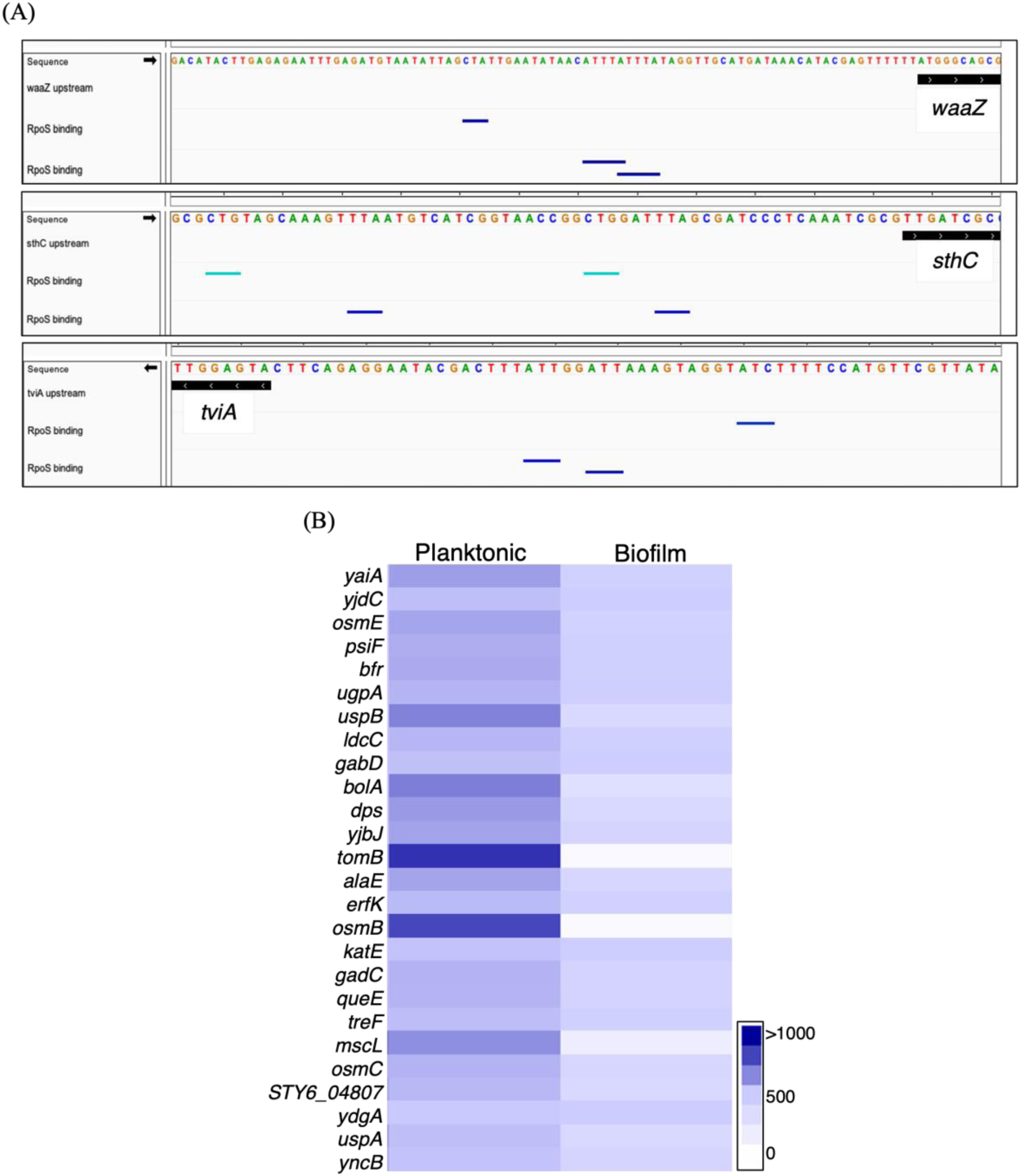
(A) Sequences of upstream regulatory regions of *waaZ*, *sthC* and *tviA* are shown as snapshots from the IGV genome browser with the RpoS binding sites, an extended -10 element followed by an AT-rich sequence, highlighted as dark blue lines (total conservation) and cyan lines (partial conservation) and (B) Hierarchical clustering of a subset of Tn-ClickSeq targets showing an enrichment of RpoS-regulated genes[59,60] in the planktonic library. For example, these include genes that confer acid resistance (*gadC*), adaptation to osmotic (*osmB*) and other environmental stresses (*psiF*, *uspA*, *uspB*), encode stress responsive regulators (*bolA* and *dps*), an iron storage protein (*bfr*), a toxin (*tomB*) and a mechanosensitive channel (*mscL*).

**Supplementary Figure 6:**
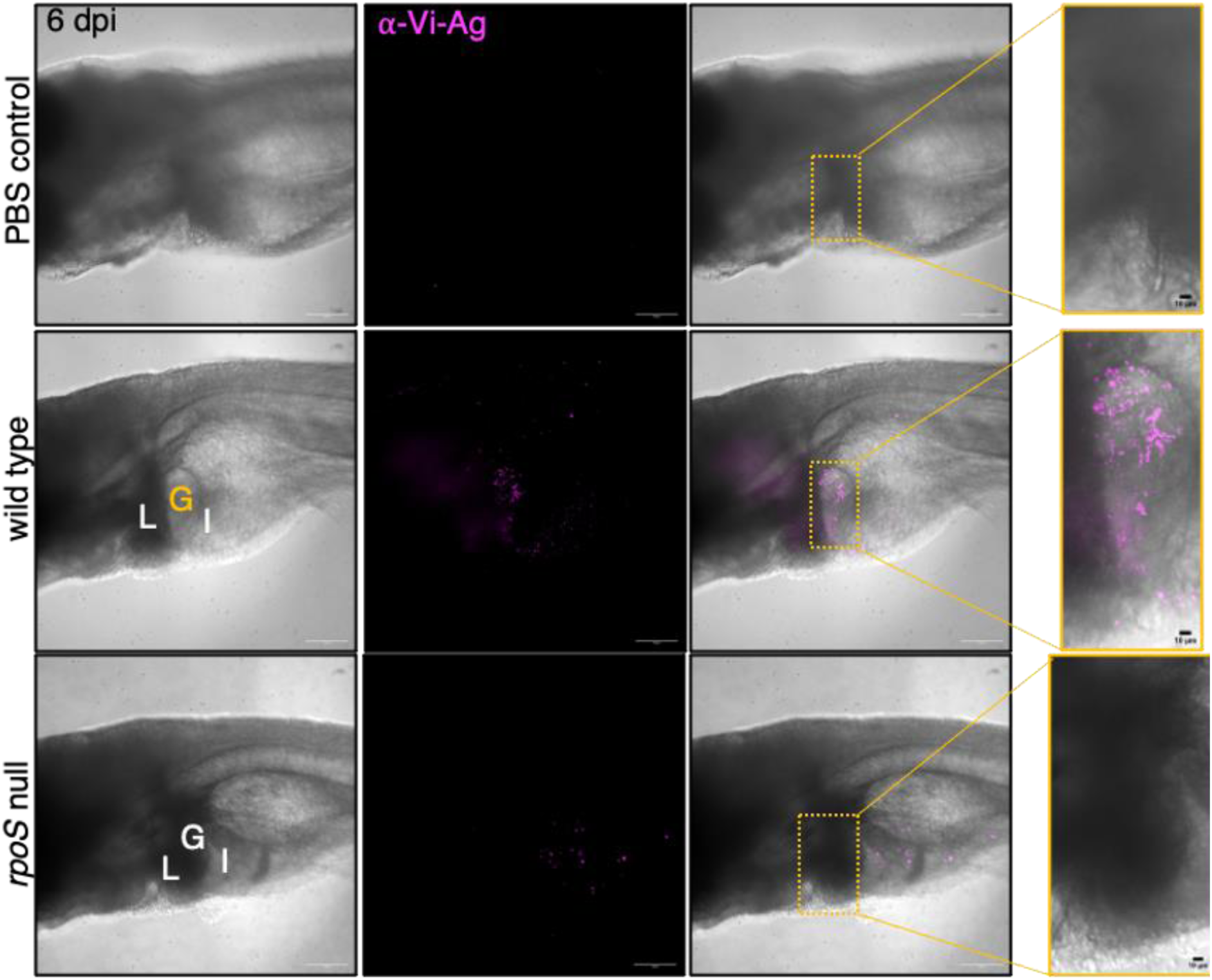
Representative whole-mount immunohistochemistry images showing the successful detection of anti-Vi-polysaccharide antibody signal (magenta) in the gall bladder region, as marked by a yellow rectangle in the merged image of a wild type infected larva at 6 dpi (middle right panel). Liver (L) and intestine (I) are marked in the bright field image on the left to pinpoint the gall bladder (G) position. The presence of Vi- antigen-positive STy clusters in the gall bladder is highlighted in the respective zoomed-in image (middle right). The gall bladder region, as marked by a yellow rectangle, of an *rpoS* null infected larva remained negative for anti-Vi-antigen antibody staining (lower panel, zoomed-in image on the right). No fluorescence was detected in the uninfected PBS control (top panel). 20X magnification, Scale bar = 100 µm for all images except for zoomed-in images with 10 µm scale bars. N = 3 with 5 to 10 larvae analyzed in each group.

**Supplementary Table 1:**
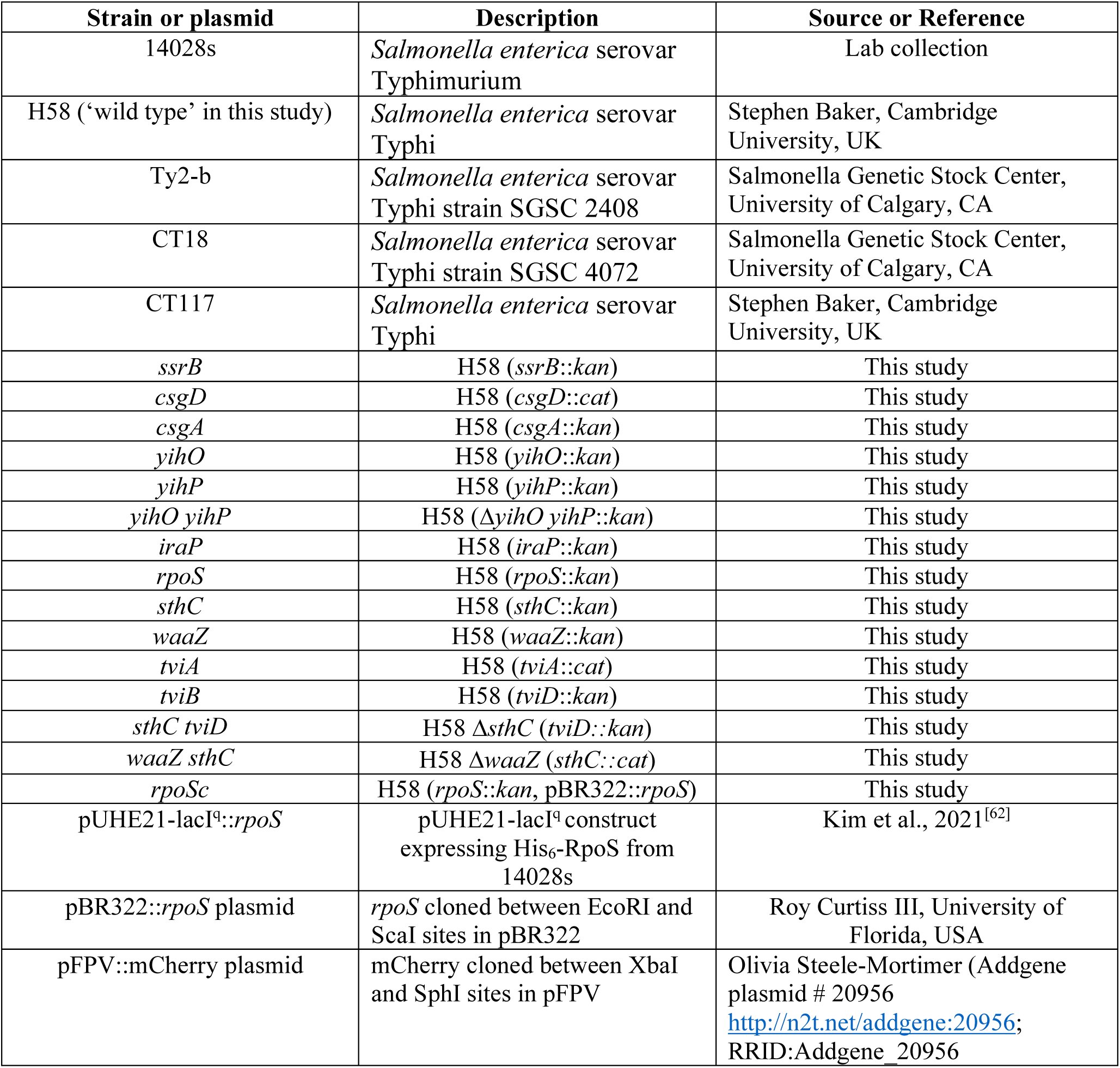
List of bacterial strains and plasmids.

**Supplementary Table 2:**
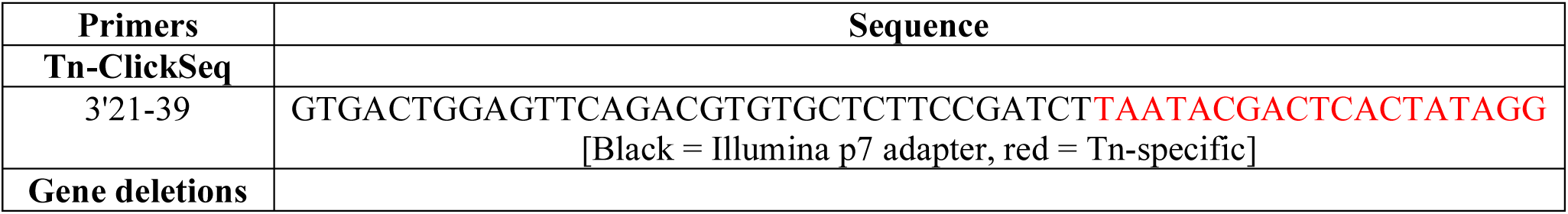
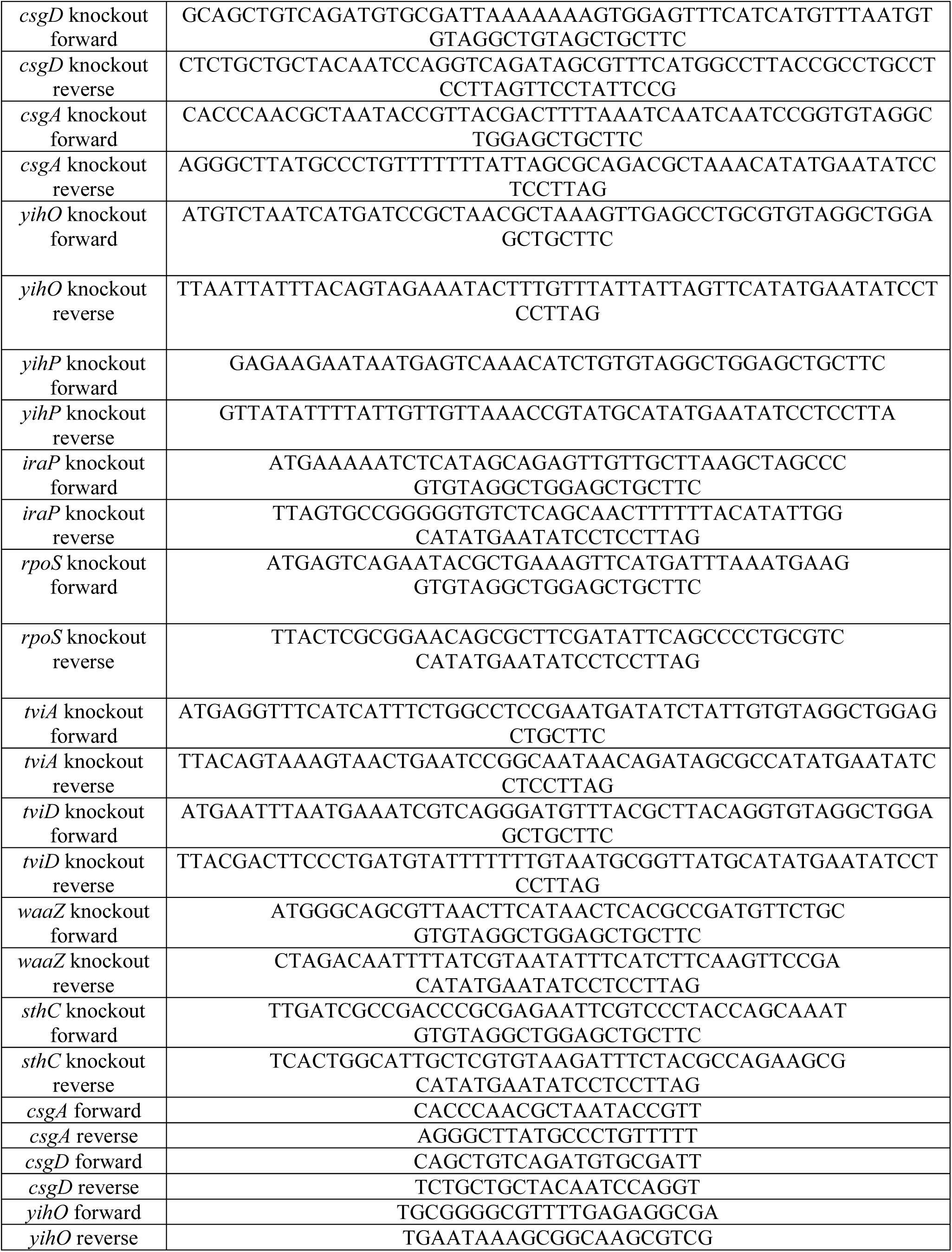
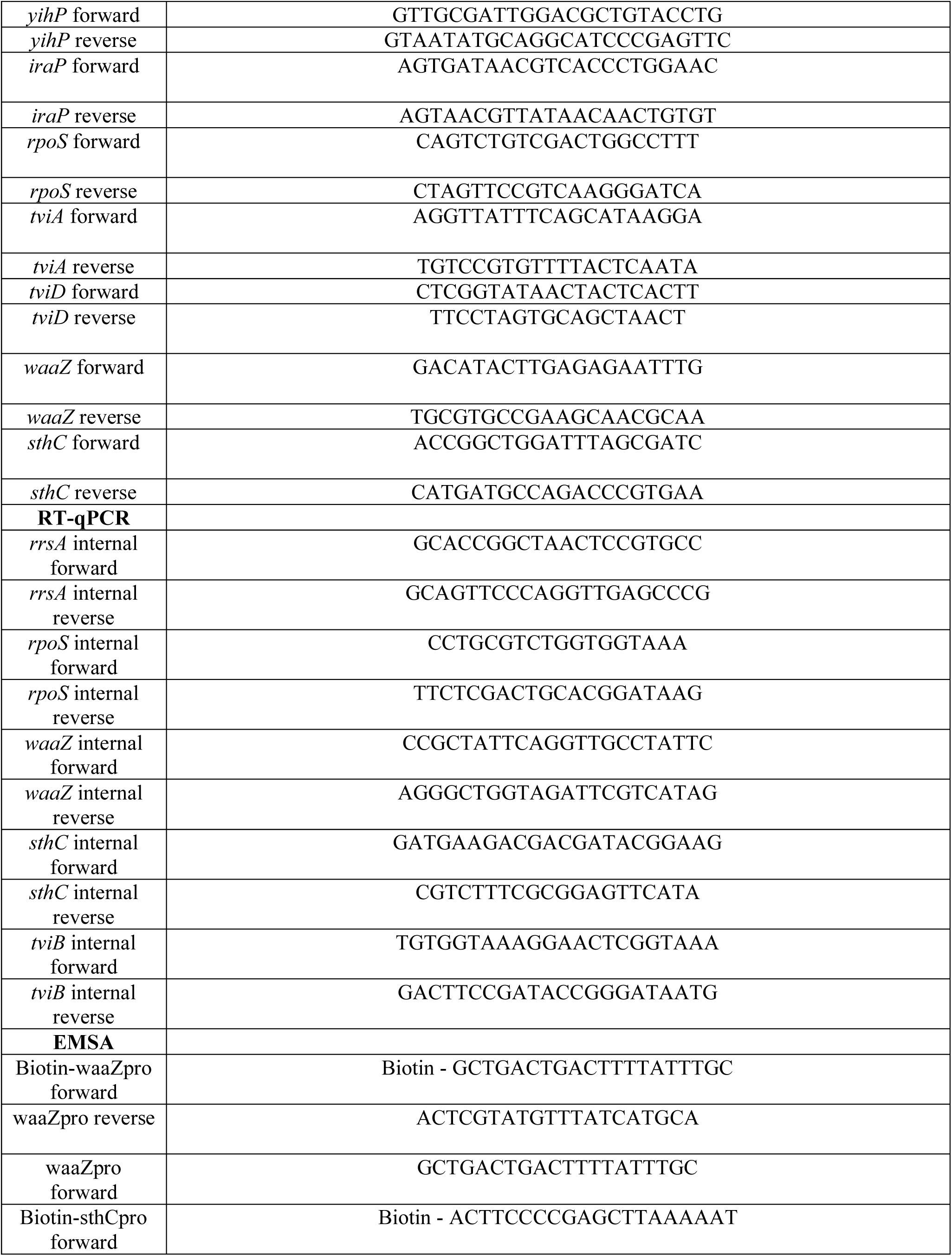
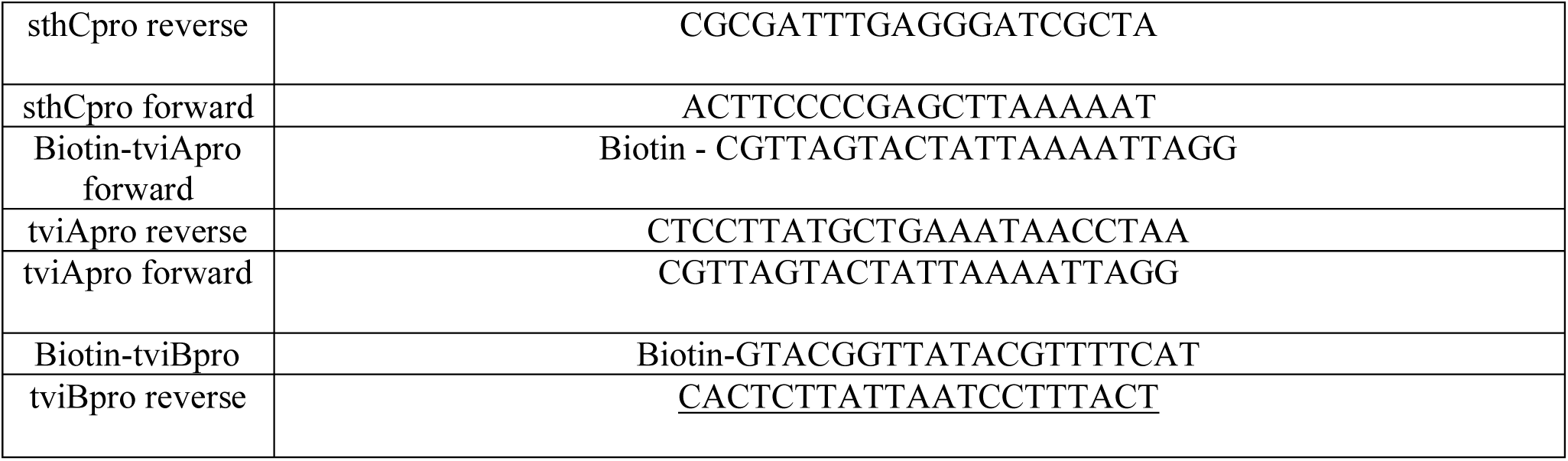
List of oligonucleotides.

